# A human electrophysiological biomarker of Fragile X Syndrome is shared in V1 of *Fmr1* KO mice and caused by loss of FMRP in cortical excitatory neurons

**DOI:** 10.1101/2025.03.19.644144

**Authors:** Sara S. Kornfeld-Sylla, Cigdem Gelegen, Jordan E. Norris, Francesca A. Chaloner, Maia Lee, Michael Khela, Maxwell J. Heinrich, Peter S. B. Finnie, Lauren E. Ethridge, Craig A. Erickson, Lauren M. Schmitt, Sam F. Cooke, Carol L. Wilkinson, Mark F. Bear

## Abstract

Predicting clinical therapeutic outcomes from preclinical animal studies remains an obstacle to developing treatments for neuropsychiatric disorders. Electrophysiological biomarkers analyzed consistently across species could bridge this divide. In humans, alpha oscillations in the resting state electroencephalogram (rsEEG) are altered in many disorders, but these disruptions have not yet been characterized in animal models. Here, we employ a uniform analytical method to show in males with fragile X syndrome (FXS) that the slowed alpha oscillations observed in adults are also present in children and in visual cortex of adult and juvenile *Fmr1^-/y^* mice. We find that alpha-like oscillations in mice reflect the differential activity of two classes of inhibitory interneurons, but the phenotype is caused by deletion of *Fmr1* specifically in cortical excitatory neurons. These results provide a framework for studying alpha oscillation disruptions across species, advance understanding of a critical rsEEG signature in the human brain and inform the cellular basis for a putative biomarker of FXS.

## Main

Fragile X syndrome (FXS) is the leading inherited cause of intellectual disability (ID) and autism spectrum disorder (ASD)^1^. In most cases, FXS is caused by transcriptional silencing of the *FMR1* gene and loss of the protein product, fragile X messenger ribonucleoprotein (FMRP). Because it is monogenic, genetically engineered animal models of FXS are available for preclinical research^2–7^. Studies in *Fmr1^-/y^* mice have revealed key aspects of neural pathophysiology and provided an opportunity to test candidate therapeutics^8–11^. To date, however, the molecules nominated to treat FXS have failed to meet prospectively defined clinical trial endpoints in humans^12^. A mismatch between preclinical promise and clinical outcomes is certainly not unique to FXS^11,12^. Given the wide differences in animal and human behavior, a bridge between the preclinical and clinical domains that measures shared alterations in circuit function is needed to better predict clinical trial outcomes^12–14^.

Field potential recordings of brain activity could be this bridge, as they are quantitative measures of pathophysiology that can be readily measured and consistently analyzed across species, a foundational requirement for translational biomarkers^14–17^. In children and adults with FXS, and in *Fmr1^-/y^* rodent models, resting-state electroencephalogram (rsEEG) recordings reveal elevated power in the gamma (30-60 Hz) frequency band (“gamma” phenotype)^18–29^. Adults with FXS also have slower alpha (8-13 Hz) oscillations^18,19,21,30^, and this shift is more reliable across individuals and testing sessions than gamma power disruptions^31–33^. However, this “alpha” phenotype has yet to be characterized in human children with FXS or in genetically engineered rodent models. A lower frequency translational biomarker of FXS would be practically important given the challenges in isolating higher frequency signals from human EEG, particularly in infants. Moreover, alpha oscillations, the most prominent rsEEG signal in humans, regulate sensory processing^34,35^ and so their disruption also fundamentally contributes to the pathology of ID^36,37^, ASD^38–40^, schizophrenia^41–44^, and other neuropsychiatric disorders^45,46^, but the disruptions have not yet been identified in preclinical models of any of these disorders. Identifying the FXS “alpha” phenotype preclinically could provide a framework for future study of the mechanistic bases of these various disruptions.

In the current study, we characterized the cross-sectional developmental trajectory of rsEEG phenotypes in male humans with FXS using spectral analysis methods that forgo traditional frequency band delineations. This approach facilitated the study of oscillations across development, and, crucially, enabled us to identify and characterize a correlate of the “alpha” phenotype in V1 of male *Fmr1^-/y^* mice. Intracortical local field potential (LFP) recordings in layer (L) 4 of V1 revealed alterations in the power and temporal dynamics of alpha-like oscillations in juvenile and adult *Fmr1^-/y^* mice, and how power alterations vary across luminance conditions. We discovered that alpha-like oscillations reflect differential activity of two genetically defined classes of cortical interneurons, parvalbumin-positive (PV+) and somatostatin-positive (SOM+) cells. Nevertheless, we found that knocking out FMRP only in cortical excitatory neurons and glia is sufficient to produce all “resting state” LFP phenotypes.

We also found that alterations in alpha-like oscillations in *Fmr1^-/y^* mice influence visual-evoked potentials (VEPs) elicited from oriented phase-reversing gratings. Intriguingly, the diminished VEPs can be improved with experience-dependent plasticity in a stimulus-specific manner. These results guide future therapeutic approaches for FXS and bridge the preclinical and clinical worlds by showing for the first time how alpha oscillation disruptions can be studied in preclinical models.

## Results

### The “alpha” phenotype is present in children with FXS, although altered physiology is not identical in children and adults

To characterize the “alpha” phenotype cross-sectionally in children and adults, rsEEG was collected using 128-channel caps in male subjects with FXS and age-matched typically developing (TD) controls across two laboratories, one which recorded from children ages 2-7 and the other which recorded from adults ages 19-44 (Fig. 1A,L). We sampled from the same electrodes across cortex for all subjects to generate an absolute power spectrum per group (FXS and TD) within each age range (Fig. 1B,M). Across all ages, we observed significantly elevated absolute power for FXS subjects below 8 Hz and above 23.5 Hz (95% confidence, non-parametric bootstrapping, n = 17 per group (p.g.) in children, n = 20 p.g. in adults).

**Figure 1.**
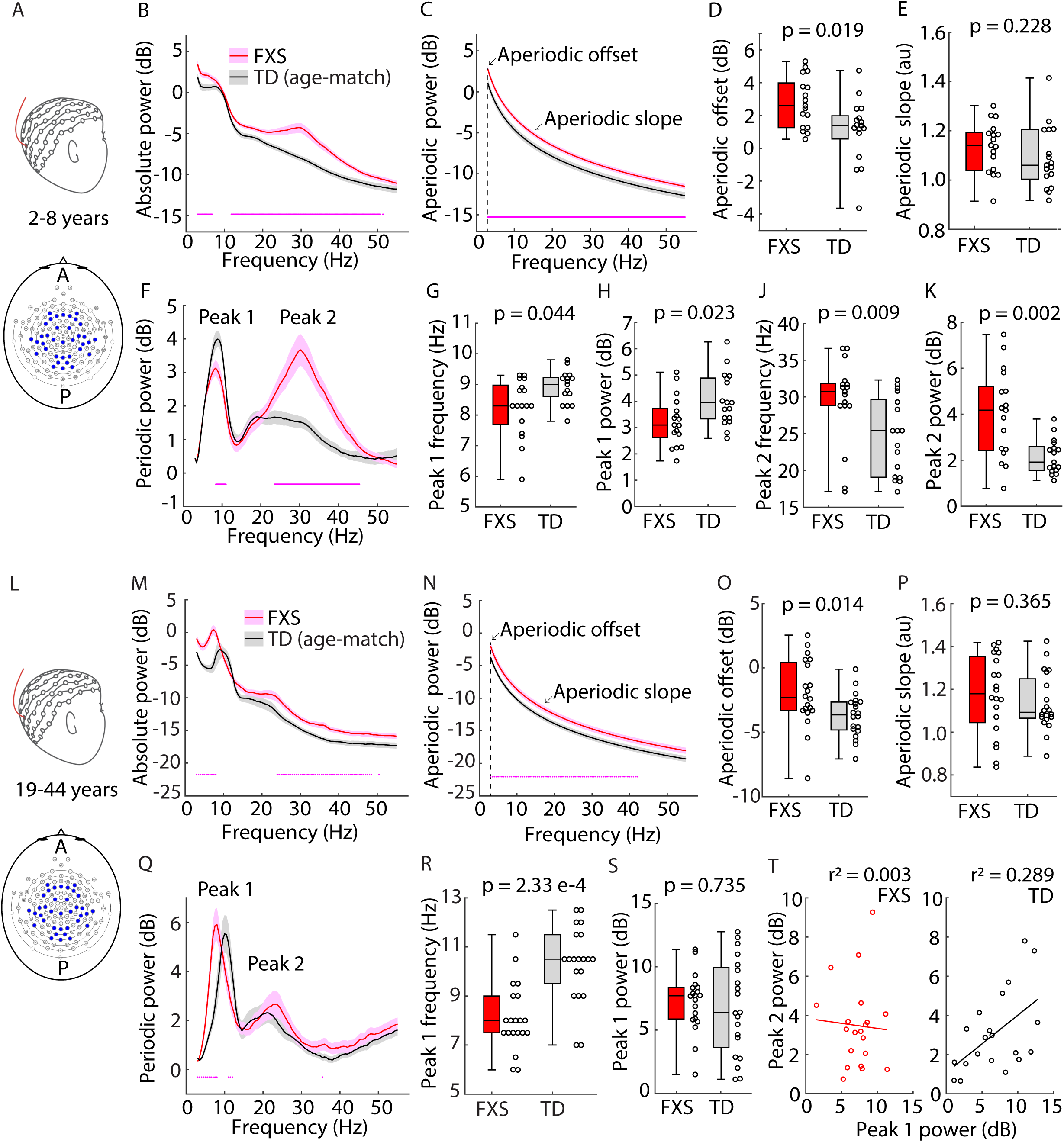
Cross-sectional developmental trajectories of rsEEG phenotypes of FXS. **A.** Experimental design. rsEEG data were collected from frontal, temporal, parietal, and occipital electrodes in FXS and TD subjects, ages 2-7 years. **B.** Absolute power spectrum (mean +/- SEM) of FXS and TD subjects (n = 17 p.g.). Dots at bottom of plots in this figure indicate points of significant difference between groups, assessed by non-parametric hierarchical bootstrap with 95% confidence. **C.** Aperiodic fit (mean +/- SEM) of the power spectra in (B). **D.** Boxplot (showing median, IQR with the box, and full range with the whiskers) and individual data points for aperiodic offset values (the power at 3 Hz, the edge of the fitting range) for FXS and TD subjects, p-value from a two-sample, two-sided Wilcoxon Rank-Sum Test (WRST, z-statistic = 2.342, effect size = 0.402). **E.** Same as (D) but for aperiodic slope (WRST z-statistic = 1.206, effect size = 0.207). **F.** Periodic power (mean +/- SEM) obtained by subtracting the aperiodic fit from the absolute spectrum for each FXS and TD subject, revealing two main oscillatory peaks. **G-K.** Boxplot and individual data points for the center frequencies of Pk1 (WRST z-statistic = - 2.017, effect size = 0.346) (G) and Pk2 (WRST z-statistic = 2.621, effect size = 0.45) (J) and the maximum power of Pk1 (WRST z-statistic = -2.273, effect size = 0.390) (H) and Pk2 (WRST z-statistic = 3.031, effect size = 0.52) (K). **L-S.** Same as A-H but for adult FXS and TD subjects, ages 19-44 years (n = 20 p.g.). For (O), WRST z-statistic = 2.448, effect size = 0.3871. For (P), WRST z-statistic = 0.906, effect size = 0.143. For (R), WRST z-statistic = -3.681, effect size = 0.582). For (S), WRST z-statistic = 0.338, effect size = 0.054. **T.** Line of best fit and linear regression r^2^ value for the correlation between the maximum power of Pk1 and Pk2 for FXS (left, effect size = 0.056) and TD adult subjects (right, effect size = 0.538).

To further analyze these spectra, we improved on existing methodology to decompose a power spectrum into aperiodic (underlying 1/f slope) and periodic (oscillatory) components (Extended Data 1A-B)^47,48^. Isolating periodic spectra improved comparisons across ages over calculating absolute power in different frequency bands, as the “alpha” rhythm sits in the theta band (3-9 Hz) in children less than 7 years old^48–50^. We found elevated aperiodic power in FXS subjects across age groups, accounting for the increased power in the absolute spectrum (Figure 1C,N). Aperiodic offset (the power at 3 Hz, the edge of the fitting range) was significantly elevated in children and adults with FXS while aperiodic slope was not significantly different between groups in either age bin (Figure 1D-E, O-P).

Removing the aperiodic component from the absolute spectra revealed that differences in periodic spectra between FXS and TD subjects were not the same for the two age groups (Figure 1F,Q). We found two periodic peaks across all groups and ages: a narrow peak one (Pk1, 3-14 Hz) and a broad, more variable peak two (Pk2, 14-38 Hz) that was strikingly aberrant in children with FXS (Extended Data C,H). Children with FXS had altered periodic power in both peaks (reduced power 8.3-11 Hz and elevated power 23.6-45.3 Hz, Fig 1F), while only Pk1 was significantly altered in adults with FXS (elevated power 3-8 Hz, reduced power 11-12 Hz from a shift, Fig 1Q).

We quantified the maximum power and center frequency of each peak for each subject. Children with FXS had a reduced maximum power of Pk1 relative to TD children (Fig. 1H) and a modest reduction in center frequency (median 8.3 Hz in FXS vs 9 Hz in TD, Fig. 1G). Pk1 was reduced in center frequency in adult FXS subjects (median 8 Hz vs. 10.5 Hz in TD adults, Fig. 1R) but Pk1 maximum power was not significantly different (Fig. 1S). In children, but not adults with FXS, the center frequency and maximum power of Pk2 were significantly increased (Fig. 1J,K; Extended Data 1K,L). The center frequency of Pk2 in children with FXS less than 60 months old clustered around the median (31.3 Hz), while the distributions for all TD and older FXS subjects were more variable (Extended Data 1E,K)^48^. Adult FXS subjects displayed a reduced correlation between the maximum power of Pk1 and Pk2 relative to TD subjects (Fig. 1T).

Overall, the most pronounced EEG phenotype in children with FXS less than 60 months old was an increase in the power and center frequency of Pk2, while the most pronounced phenotype of adults with FXS was the shift in center frequency of Pk1; both, along with the increased aperiodic power in FXS subject of all ages, were highly consistent across cortical regions (Extended Data 2 and 3). The “gamma” phenotype was due to elevated aperiodic power in adults, but due to elevated periodic *and* aperiodic power in children. Critically, both children and adults with FXS displayed the “alpha” phenotype (the shift in center frequency of Pk1), although it was more pronounced in adults. For children with FXS, Pk1 was reduced both in maximum power and center frequency.

### A correlate of the “alpha” phenotype is present in V1 of adult *Fmr1^-/y^* mice

We next used our analyses to investigate if mice also have a periodic Pk1, and if so, whether its center frequency is reduced in *Fmr1^-/y^* mice. Adult *Fmr1^-/y^* and littermate wild-type (WT) mice (postnatal day (p)70-115, n = 11 p.g.) were implanted with screw-type electrodes in the skull over primary somatosensory cortex (S1) and V1. (Fig. 2A). One week later, the awake, freely moving mice were sequentially placed in a video-monitored chamber while EEG data was continuously recorded. Behavioral scoring using video recordings facilitated extraction of 100 sec of “resting-state” (i.e., awake but stationary) data from each animal.

**Figure 2.**
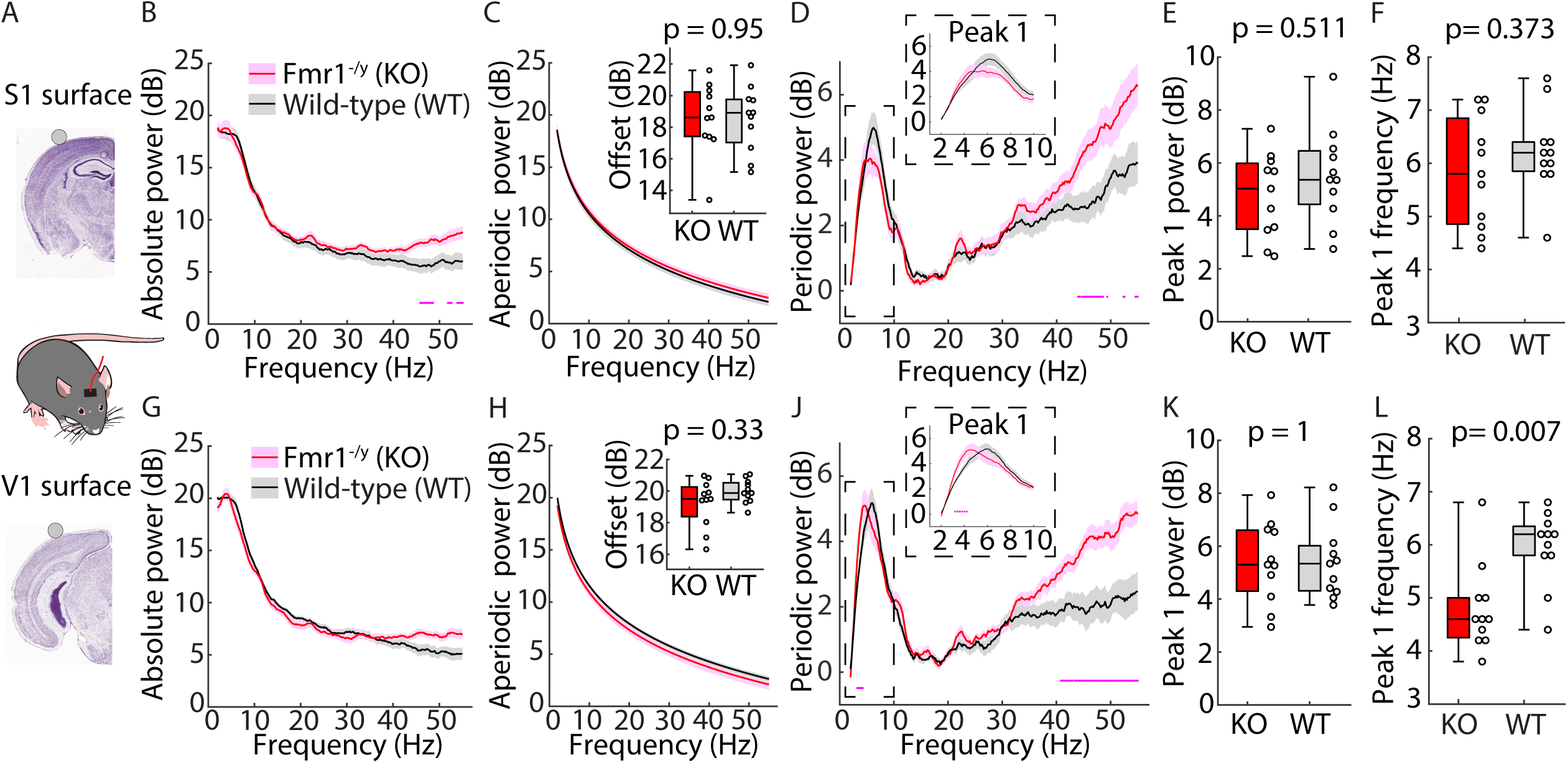
A correlate of the “alpha” phenotype is present in V1 of adult *Fmr1^-/y^* mice. **A.** Experimental design. EEG data were collected from electrodes on the cortical surface S1 and V1 of freely-moving adult *Fmr1^-/y^* (KO) and littermate WT mice (p70-115). 100 sec of data while the mice were awake but stationary was analyzed as a comparable “resting-state” dataset. **B.** Absolute power spectrum (mean +/- SEM) from the S1 electrode in KO and WT mice (n = 11 p.g.). Dots at bottom of plots in this figure indicate points of significant difference between groups, assessed by non-parametric hierarchical bootstrap with 99% confidence. **C.** Aperiodic fit (mean +/- SEM) of the power spectra in (B). Inset: Boxplot (median, IQR, and full range) and individual data points for aperiodic offset values (the power at 2 Hz) for *Fmr1* KO and WT mice, WRST (z-statistic = 0.066, effect size = 0.014). **D.** Periodic power for the S1 electrode (mean +/- SEM). **E-F.** Boxplot and individual data points for the maximum power of Pk1 (WRST z-statistic = -0.657, effect size = 0.14) (E) and the center frequency of Pk1 (WRST z-statistic = -0.89, effect size = 0.19) (F). **G-L**. Same as B-F but for the V1 electrode. For (H), WRST z-statistic = -0.985, effect size = 0.21. For (K), WRST z-statistic and effect size both = 0. For (L), WRST z-statistic = -2.705, effect size = 0.577.

We observed significantly elevated absolute power from S1 electrodes in *Fmr1^-/y^* mice between 45.6-55 Hz (99% confidence, non-parametric bootstrapping) and a similar trend in V1 (Fig. 2B,G), replicating reports of elevated gamma power in *Fmr1^-/y^*mice^23–25,28,29^. We next analyzed the aperiodic and periodic components. Unlike in humans, there was no difference in aperiodic power or offset between the two groups (Fig. 2C,H). Instead, both genotypes had a broad increase in periodic power at high frequencies which was larger in *Fmr1^-/y^*mice (significant difference 44.8-55 Hz for S1, 41.2-55 Hz for V1, Figure 2D,J). There was a robust periodic Pk1 between 2-13 Hz in both genotypes and no significant difference between genotypes in the maximum power of Pk1 in either S1 or V1 (Fig. 2E,K). Using the electrode over V1, we found a significant reduction in the center frequency of Pk1 in *Fmr1^-/y^* mice (median 4.6 Hz) relative to WT mice (median 6.2 Hz) (Fig. 2L). This shift in center frequency of Pk1 in V1 of adult *Fmr1^-/y^* mice was consistent with the shift in Pk1 seen in adult humans with FXS: we identified a correlate of the “alpha” phenotype in mice.

### Intracortical recordings in V1 recapitulate the “alpha” phenotype in juvenile and adult mice and reveal a two subpeak structure to periodic Pk1

Our discovery of the parallel Pk1 phenotype in *Fmr1^-/y^*mice supports work positing a commonality between theta oscillations in V1 of mice and human alpha oscillations^51,52^. 3-6 Hz alpha-like oscillations in mouse V1 are present in all cortical layers but strongest in L4^51^. We further characterized the “alpha” phenotype by implanting *Fmr1^-/y^* and littermate WT mice with LFP microelectrodes targeted to L4 of V1. After recovery, the mice were habituated to the recording setup (head-fixation in front of a monitor). Head-fixation permitted control of visual input and limited motion artifacts in high frequencies of the LFP. After two days of habituation, we characterized 150 sec of “resting-state” data while the monitor was turned off (mice were in the dark), analyzing data from juvenile and adult mice (p30-150, n = 44 p.g.).

We observed a significant elevation in absolute power in *Fmr1^-/y^* mice between 26-96.6 Hz (Fig. 3A). We found no difference in the aperiodic fit between genotypes (Extended Data 4A-F). Removing the aperiodic component from the absolute spectra revealed the full broad shape of the periodic high frequency signal in both genotypes and a significant elevation in this periodic power in *Fmr1^-/y^*mice between 22-133.4 Hz (Fig. 3B). Intriguingly, periodic Pk1 (2-10 Hz) was composed of two smaller sub-peaks (Pk1a and Pk1b). There was a significant difference in periodic power between genotypes from 4.2-5.4 Hz, within Pk1a. Both the center frequency and maximum power of Pk1a (but not Pk1b) were reduced in *Fmr1^-/y^*mice (Fig. 3C-D). The intracortical “alpha” phenotype was contained within Pk1a and was accompanied by a power reduction.

**Figure 3.**
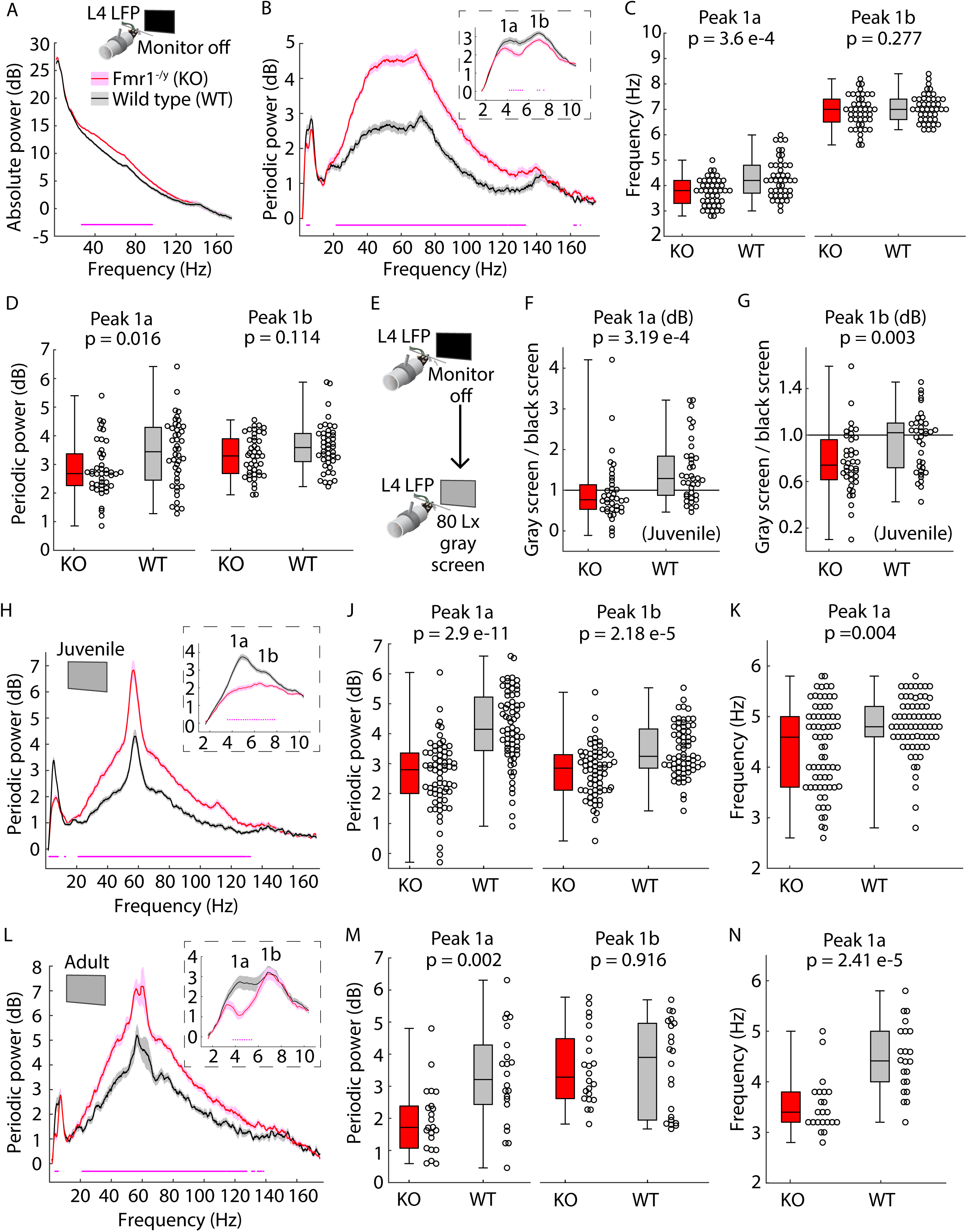
Genotype differences in periodic Pk1 in V1 change across luminance conditions differentially for juvenile and adult mice. **A.** Inset: Experimental design. LFP data were collected from L4 of V1 in KO and littermate WT mice (p30-150) head-fixed in front of a monitor. Mice were habituated to head-fixation for two days before data collection. Main: Absolute power spectrum (mean +/- SEM) from the L4 LFP electrode in V1 of KO and WT mice (n = 44 p.g.) while the monitor was turned off (i.e., the mice were in the dark). Dots at bottom of plots in this figure indicate points of significant difference between groups, assessed by non-parametric hierarchical bootstrap with 99% confidence. **B.** Periodic power (mean +/- SEM) for KO and WT mice. Periodic Pk1 is composed of two sub-peaks (1a and 1b). **C.** Boxplot (median, IQR, and full range) and individual data points for the center frequency of Pk1a and Pk1b. WRST z-statistic = -3.57, effect size = 0.381 (Pk1a) and WRST z-statistic = -1.087, effect size = 0.1158 (Pk1b). **D.** Same as (C) but for the maximum power of Pk1a and Pk1b. WRST z-statistic = -2.408, effect size = 0.257 (Pk1a) and WRST z-statistic = -1.581, effect size = 0.169 (Pk1b). **E.** Next, the monitor was turned on to display a static, iso-luminant gray screen (80 Lx). **F.** Ratio of black screen and gray screen Pk1a maximum power for juvenile KO and littermate WT mice, WRST z-statistic = -3.6, effect size = 0.413. **G.** Same as (F) but for Pk1b periodic power, WRST z-statistic = -2.997, effect size = 0.344. **H.** Periodic power spectrum (mean +/- SEM) from the L4 LFP electrode in V1 of juvenile KO and littermate WT mice (p30-40) during gray screen (n = 67 p.g.). **J.** Boxplot and individual data points for the maximum power of Pk1a and Pk1b. WRST z-statistic = -6.653, effect size = 0.575 (Pk1a) and WRST z-statistic -4.245, effect size = 0.367 (Pk1b). **K.** Boxplot and individual data points for the center frequency of Pk1a, WRST z-statistic = -2.878, effect size = 0.249. **L-N.** Same as (H-K) but for adult KO and littermate WT mice (p70-150) during gray screen (n = 22 p.g.). For (M), WRST z-statistic = -3.181, effect size = 0.48 (Pk1a) and WRST z-statistic = 0.106, effect size = 0.016 (Pk1b). For (N), WRST z-statistic = - 4.223, effect size = 0.637.

### Periodic Pk1 phenotypes change across luminance conditions differentially in juvenile and adult *Fmr1^-/y^* mice

In WT mice, alpha-like oscillations are strongly driven by a flash of light^51^, so we investigated how luminance affects “resting-state” periodic Pk1 phenotypes by displaying a static, isoluminant gray screen (Fig. 3E). Across all animals, luminance increased aperiodic power below 60 Hz and significantly increased aperiodic offset and slope (Extended Data 4G-H). Intriguingly, for periodic power, luminance revealed developmental differences, so we separately analyzed juvenile (p30-40) and adult (p70-150) mice. In juvenile mice (n = 38 p.g.), luminance enhanced the maximum power of Pk1a in WT, but not in *Fmr1^-/y^*mice, and it suppressed the maximum power of Pk1b in *Fmr1^-/y^*, but not WT mice (Extended Data 4J), thereby exacerbating genotype differences (Fig. 3F-G). Luminance did not have the same effect on Pk1b in adult *Fmr1^-/y^* mice (Extended Data 4K-L).

We next analyzed periodic power spectra for gray screen “resting-state” data. Luminance induced a narrowband periodic signal around 60 Hz in both genotypes^53^. This signal sat atop the broad high-frequency periodic signal seen before, and for both juvenile (n = 67 p.g.) and adult mice (n = 22 p.g.), periodic power was significantly elevated in *Fmr1^-/y^* mice between 21.2-127.6 Hz (Fig. 3H,L). For juvenile *Fmr1^-/y^* mice, periodic power was significantly reduced between 3.6-7.8 Hz; for adult mice the reduction was significant between 3.8-5.4 Hz. Luminance induced a significant genotype difference in the maximum power of Pk1a and Pk1b for juvenile mice, but only in Pk1a for adult mice (Fig. 3J,M).

In adult *Fmr1^-/y^* mice, the center frequency of Pk1a was more strongly shifted relative to WT than in juveniles (Fig. 3K,N). For gray screen data, the center frequency of Pk1a was significantly smaller in adult relative to juvenile *Fmr1^-/y^* mice (Extended Data 4N). In juvenile animals, the maximum power of Pk1a and Pk1b and Pk1a center frequency separated the two genotypes’ distributions, while in adults, the two genotypes were separated by Pk1a frequency and power (Extended Data 4O,P). This observation somewhat parallels the shift from power to center frequency of Pk1 more strongly separating the distributions of human subjects across development (Extended Data 1D,J).

Previous studies have shown a relationship between the phase of mouse alpha-like oscillations in L4 of V1 and the amplitude of oscillations within the frequency range of the human Pk2^51,54^. We have reproduced that finding in WT mice and shown that this phase-amplitude coupling is impaired and phase-shifted in *Fmr1^-/y^* mice (Extended Data 5), echoing the reduced correlation between the power of periodic Pk1 and Pk2 in humans with FXS (Figure 1T, Extended Data 1M).

### Altered temporal dynamics of alpha-like oscillations in *Fmr1^-/y^* mice

In humans, alpha oscillations occur in bursts and burst dynamics are altered in FXS^55^. To test if dynamics are similarly altered in *Fmr1^-/y^* mice, we bandpass-filtered 100 sec of continuous “resting-state” LFP data between 2-10 Hz (periodic Pk1) and used thresholding to identify “microbursts” (Fig. 4A). In *Fmr1^-/y^* mice, microbursts occurred more frequently but lasted for shorter durations across both ‘monitor off’ and ‘gray screen’ data (Fig. 4B,D). Microbursts tended to occur during periods of behavioral quiescence, but there was no evidence of hyperactivity in *Fmr1^-/y^* mice (Fig. 4C,E)^23,56^. Altered microburst dynamics were present in both juvenile and adult mice (Extended Data 6A-D) and were detectable in L4 but not at the cortical surface (Extended Data 6E). Similarly, we could not detect these effects in our rsEEG data from humans, largely due to immense variability in TD data (Extended Data 6G). However, previous work identified elevated burst counts in FXS male subjects in inter-stimulus intervals of a task (not resting-state data)^55^, consistent with our results in *Fmr1^-/y^*mice. For resting-state data, intracortical recordings revealed differences in temporal dynamics not detectable from the surface.

**Figure 4.**
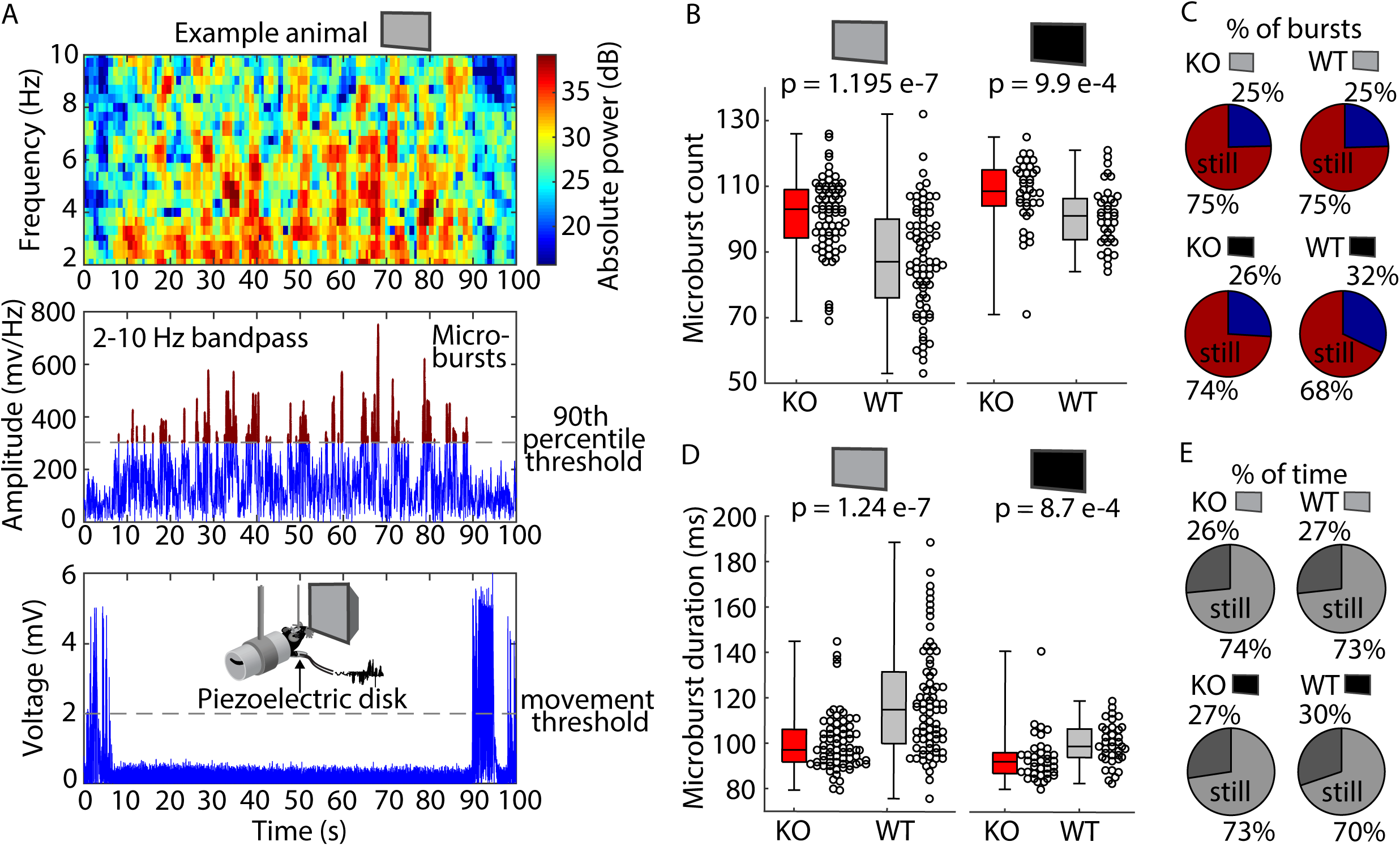
Alpha-like oscillations in V1 of *Fmr1^-/y^* mice exhibit altered temporal dynamics. **A.** Methodology. Time series from an example animal viewing a static gray screen. Across 100 sec of continuous data, power between 2-10 Hz fluctuates in a burst-like pattern. Band-passing the 2-10 Hz signal revealed the number of times the signal crosses the 90^th^ percentile band-passed value (microburst count) and the length of time the signal stays above the 90^th^ percentile threshold after each crossing (microburst duration). Timing of bursts was compared to timing of movement bouts, measured through a piezoelectric disk under the animal’s forepaw. Altered pressure on the disk during movement was recorded as voltage deflection, and voltages values above 2 mV constituted movement. **B.** Boxplot (median, IQR, and full range) and individual data points for the microburst count for gray screen (left) and black screen (right). WRST z-statistic = 5.102, effect size = 0.424 (gray screen) and WRST z-statistic = 3.293, effect size = 0.38 (black screen). **C.** Distribution of average burst activity across all KO (left) and WT (right) animals during gray screen (top) and black screen (bottom) during moving and still states. **D.** Same as (B) but for microburst duration. WRST z-statistic = -5.098, effect size = 0.423 (gray screen) and WRST z-statistic = -3.3, effect size = 0.385 (black screen). **E.** Percentage of time spent moving and still across all KO (left) and WT (right) animals during gray screen (top) and black screen (bottom).

### Alpha-like oscillations in mice are linked to differential activity of genetically defined classes of inhibitory interneurons

Thus far, we have characterized the “alpha” phenotype in both children and adults with FXS and established its correlate in V1 of *Fmr1^-/y^*mice through surface and intracortical recordings. We next investigated the mechanistic bases for the alterations observed in periodic Pk1. In mice, PV+ interneurons coordinate cortical theta and gamma oscillations (i.e., frequencies within periodic Pk1 and the broad high frequency periodic peak)^57–59^, and PV+ interneuron density and activity is deficient in FXS in both mice and humans^60,61^. Alpha oscillations are broadly tied to cortical inhibition^35,62–64^ and reflect activity in the thalamocortical loop^34,52,65^, but whether they also reflect activity of inhibitory interneurons is unclear. We investigated if inhibiting PV+ neurons in V1 of WT mice would mimic the changes to the L4 LFP seen in juvenile *Fmr1^-/y^* mice, particularly within periodic Pk1a and Pk1b.

We re-analyzed previously published data wherein chemogenetic methods were used to temporarily inactivate PV+ cells in V1^66^. We analyzed 150 sec of LFP data where mice (n = 16) expressed the chemogenetic actuator hM4Di in PV+ neurons of V1 and viewed a static gray screen before and after systemic injection of Clozapine N-Oxide (CNO) to trigger the hM4Di receptor. Inactivating PV+ cells increased absolute power between 1.6-110.4 Hz (Fig. 5A). This effect resulted from an increase in aperiodic power between 1.6-106.2 Hz (as seen in the human FXS rsEEG) and an increase in periodic power between 9.6-115.8 Hz (as seen in the *Fmr1^-/y^* mouse LFP) (Fig. 5B-D). PV+ neuronal inactivation eliminated periodic Pk1b power (significant difference 6.8-8.2 Hz) but did not affect Pk1a power or center frequency (Fig. 5C,E-F and Extended Data 7E). It also increased the frequency and shortened the duration of Pk1 microbursts without changing overall movement (Fig. 5G-J). Since this manipulation failed to affect periodic Pk1a, additional mechanisms may underlie the alterations to periodic Pk1.

**Figure 5.**
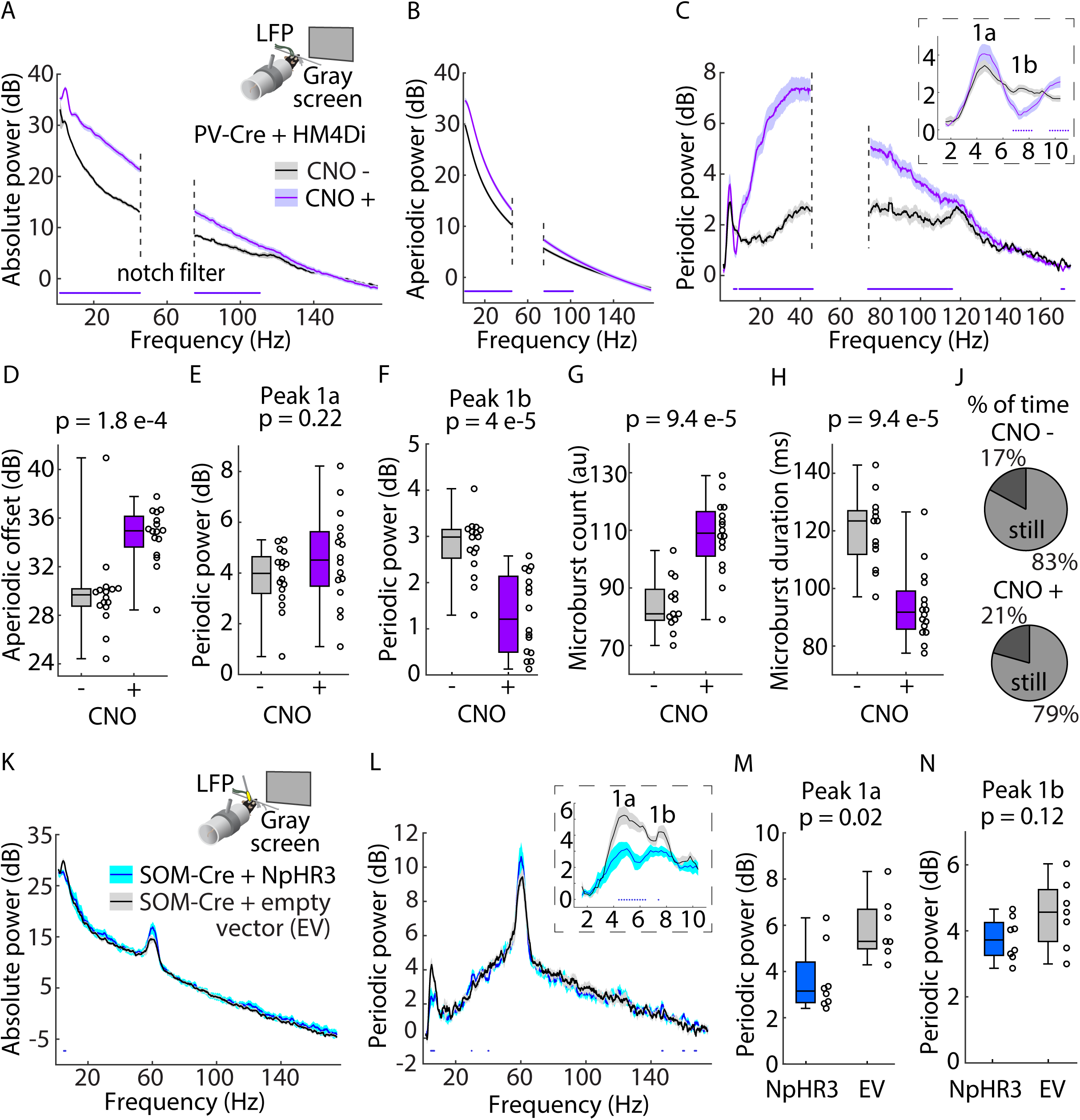
Inhibiting PV+ or SOM+ interneurons in WT mice partially recapitulates the changes to the V1 L4 LFP seen in *Fmr1* KO mice. **A.** Inset: experimental design. LFP data were collected from L4 of V1 in PV-Cre mice (p50-80) with chemogenetic actuator hM4Di expressed in V1^66^. The mice were head-fixed in front of a monitor displaying a static, iso-luminant gray screen. Main: Absolute power spectrum (mean +/- SEM) from the L4 LFP electrode in V1 of PV-Cre mice (n = 16) before and after systemic injection of CNO to activate the hM4Di receptor. Frequencies between 45-75 Hz are not analyzed due to a notch filter. Dots at bottom of plots in this figure indicate points of significant difference between groups, assessed by non-parametric hierarchical bootstrap with 99% confidence. **B.** Aperiodic fit (mean +/- SEM) of the power spectra in (A). **C.** Periodic power spectrum (mean +/- SEM). **D.** Boxplot (median, IQR, and full range) and individual data points for aperiodic offset values (the power at 1.5 Hz), WRST z-statistic = -3.75, effect size = 0.6629. **E-F.** Boxplot and individual data points for the maximum power of Pk1a (E) and Pk1b (F). WRST z-statistic = -1.225, effect size = 0.217 (E) and WRST z-statistic = 4.127, effect size = 0.73 (F). **G-H.** Same as (E-F) but for microburst count (G) and microburst duration (H). WRST z-statistic = -3.903 (count) and 3.903 (duration), effect size = 0.738 for both. **J.** Percentage of time spent moving and still across all PV-Cre mice before (top) and after (bottom) CNO injection. **K.** Inset: Same as (A) but for SOM-Cre mice (p50-70) with either an inhibitory opsin virus (NpHR3) or an empty control vector (EV) expressed in V1. Main: Absolute power spectrum (mean +/- SEM) from the L4 LFP electrode in V1 of SOM-Cre+NpHr3 (n = 8) and SOM-Cre+EV mice (n = 7) during ‘laser on’ periods. **L.** Sane as (C) but for SOM-Cre mice. **M-N.** Same as (E-F) but for SOM-Cre mice. Due to the smaller sample size, z-statistics and effect sizes not reported.

SOM+ interneurons also regulate cortical oscillations^58,59,67^ and inhibiting SOM+ cells diminishes visually-evoked theta oscillations^54^. To examine if altered activity of SOM+ interneurons affects Pk1a, we used optogenetics to temporarily inactivate SOM+ activity in V1 while recording “resting-state” LFP during presentation of a static gray screen (Fig. 5K).

Optogenetic inactivation reduced absolute power between 4.6-6.2 Hz in mice expressing the inhibitory actuator Halorhodopsin (NpHR3) in SOM+ neurons of V1 (n = 8), relative to control mice (n = 7). The manipulation had no effect on aperiodic power (Extended Data 7F-J), but periodic power of Pk1a (but not Pk1b) was reduced (significant difference between 4.4-6.4 Hz) (Fig. 5L-N). The manipulation did not affect the center frequency of either subpeak (Extended Data 7K). The power of Pk1a and Pk1b are therefore separately affected by inhibition of SOM+ and PV+ cells, respectively. Deficient activity of multiple classes of interneurons might contribute to the electrophysiological differences seen in the LFP of *Fmr1^-/y^*mice.

### Loss of FMRP in cortical excitatory neurons and glia in mice causes the “alpha” phenotype

Given the relationship between periodic Pk1 power and the activity of inhibitory interneurons, we then examined if sparing FMRP expression in inhibitory interneurons might prevent the altered LFP of *Fmr1^-/y^* mice. We employed a genetic strategy to selectively knockout *Fmr1* from excitatory neurons and glia in cortex (EMX1-*Fmr1* KO), sparing its expression in cortical inhibitory interneurons and subcortically^68^. We used juvenile mice (p30-40) and implanted electrodes in L4 of V1 to record “resting-state” LFP while the monitor was off or displaying a static gray screen. WT mice in this triple transgenic line (n = 20) showed no difference from WT mice used previously (n = 9) (Extended Data 8A-F). As shown in Figure 6 and Extended Data 8G-M, all the resting-state LFP phenotypes replicated in EMX1-*Fmr1* KO mice (n = 15). Altered physiology, including the “alpha” phenotype, is caused by loss of FMRP in excitatory neurons.

**Figure 6.**
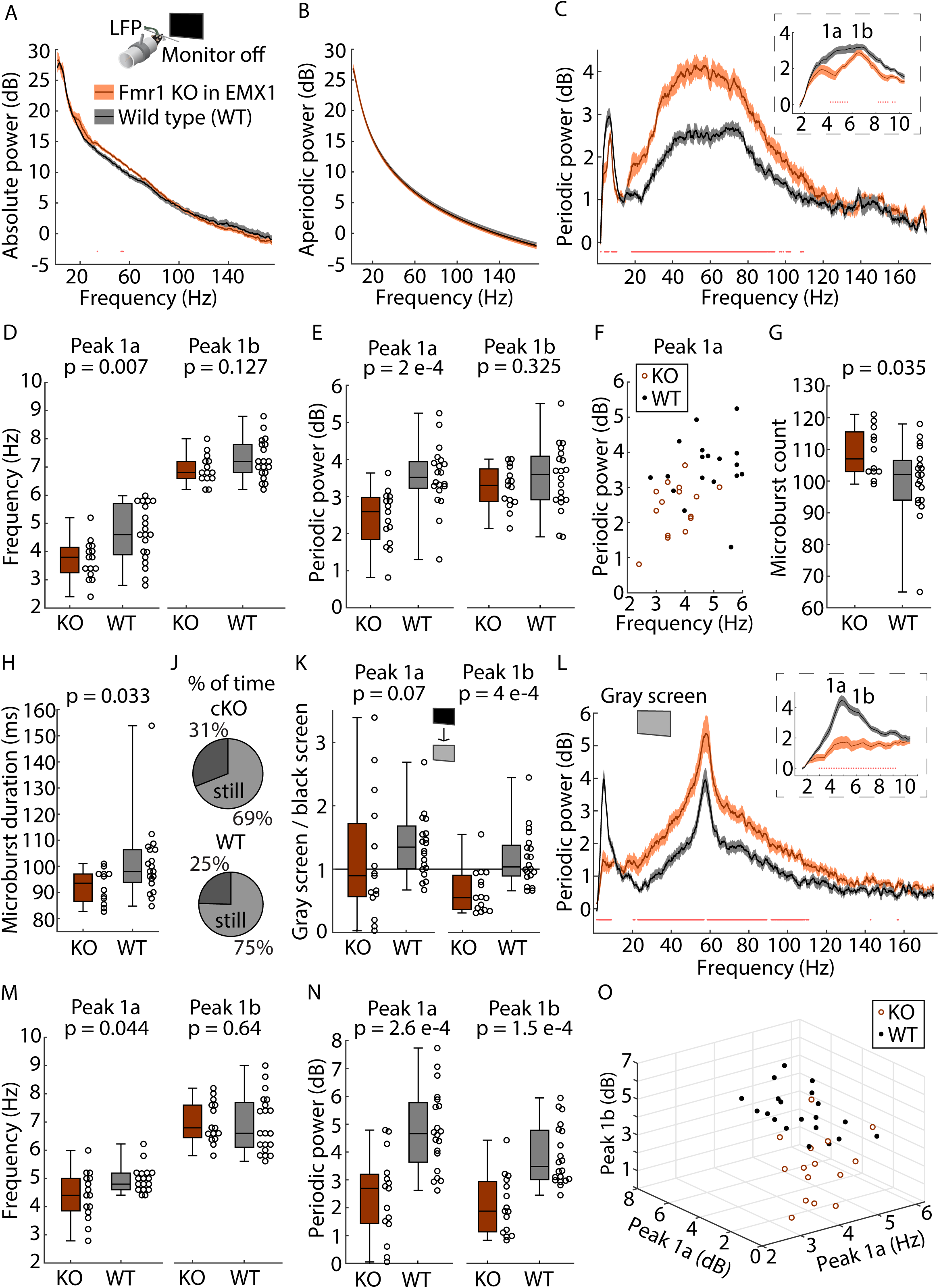
Loss of FMRP in cortical excitatory neurons and glia causes the changes in the V1 L4 LFP seen in *Fmr1* KO mice. **A.** Inset: Experimental design. LFP data were collected from L4 of V1 in juvenile (p30-40) WT mice and mice with *Fmr1* specifically knocked out of cortical excitatory neurons and glia expressing the EMX1 promotor (EMX1-*Fmr1* KO). Mice were head- fixed in front of a monitor and habituated to head-fixation for two days before data collection. Main: Absolute power spectrum (mean +/- SEM) from the L4 LFP electrode in V1 of EMX1-*Fmr1* KO (Cre+/Fmr1-, n = 15) and WT mice (Cre+/Fmr1+ (Cre/WT), n = 10 and Cre-/Fmr1-(WT/Fmr1 KO), n = 10) while the monitor was turned off (i.e., the mice were in the dark). Dots at bottom of plot in this figure indicate points of significant difference between groups, assessed by non-parametric hierarchical bootstrap with 99% confidence. **B.** Aperiodic fit (mean +/- SEM) of the power spectra in (A). **C.** Periodic power spectrum (mean +/- SEM) for EMX1-*Fmr1* KO and WT mice. **D-E.** Boxplot (median, IQR, and full range) and individual data points for the center frequency (D) and maximum power (E) of Pk1a and Pk1b. In (D), WRST z-statistic = - 2.69, effect size = 0.455 (Pk1a) and WRST z-statistic = -1.527, effect size = 0.258 (Pk1b). In (E), WRST z-statistic = -3.717, effect size = 0.628 (Pk1a) and WRST z-statistic = -0.983, effect size = 0.166 (Pk1b). **F.** Separation of genotypes by Pk1a center frequency and maximum power during monitor off (black screen). **G-H.** Same as (D-E) but for microburst count (G) and microburst duration (H). For (G), WRST z-statistic = 2.112, effect size = 0.379. For (H), WRST z-statistic = -2.13, effect size = 0.383. **J.** Percentage of time spent moving and still across all EMX1-*Fmr1* KO mice (top) and WT mice (bottom). **K.** Ratio of black screen and gray screen periodic Pk1a and Pk1b maximum power values for each mouse. WRST z-statistic = -1.783, effect size = 0.301 (Pk1a) and WRST z-statistic = -3.55, effect size = 0.6 (Pk1b). **L.** Same as (A) but while the monitor was turned on and displaying an iso-luminant gray screen. **M-N.** Same as (D-E) but for gray screen. In (M), WRST z-statistic = -2.01, effect size = 0.34 (Pk1a) and WRST z-statistic = 0.467, effect size = 0.079 (Pk1b). In (N), WRST z-statistic = -3.65, effect size = 0.617 (Pk1a) and WRST z-statistic = -3.78, effect size = 0.64 (Pk1b). **O.** Same as (F) but during gray screen and considering the added dimension of Pk1b maximum power.

### Periodic Pk1 phenotypes predict altered visual-evoked activity in *Fmr1^-/y^* mice

Finally, we consider how the alterations in periodic Pk1 in *Fmr1^-/y^*mice might affect VEPs. In humans, the phase of alpha oscillations at stimulus onset affects the amplitude of the subsequent VEP, and VEPs are enhanced to rhythmic stimuli within the alpha frequency range^69–72^. We hypothesized a conserved relationship between periodic Pk1 oscillations and VEPs and predicted altered onset responses and VEPs to rhythmic stimuli in *Fmr1^-/y^* mice.

After collecting “resting-state” data from *Fmr1^-/y^* and littermate WT mice implanted with LFP electrodes in L4 of V1, we presented a grating stimulus at a fixed orientation which phase-reversed at frequencies ranging from 2-15 Hz, with 10 sec of gray screen between each frequency (Fig. 7A). We separately analyzed the stimulus onset response (first 250 msec), which was stereotyped across temporal frequencies, and the subsequent “entrainment” period for each frequency (Fig. 7B, Extended Data 9D). In both WT and *Fmr1^-/y^* mice (n = 37 p.g.), the amplitude of the first negativity (N1) of the onset response depended on the phase of periodic

**Figure 7.**
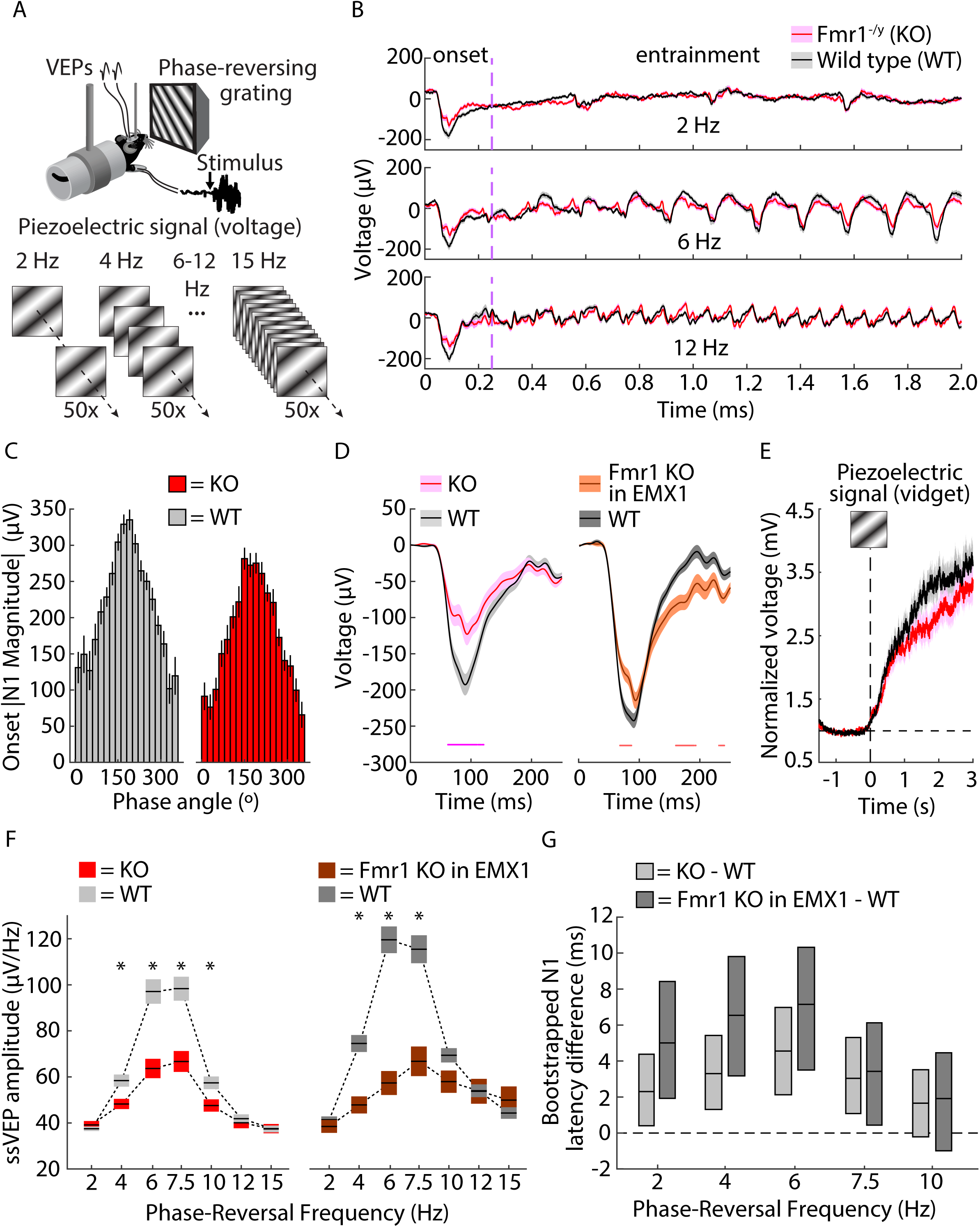
VEPs in *Fmr1* KO mice are either reduced in magnitude, delayed, or both, depending on the stimulus. **A.** Experimental design. LFP data from L4 of V1 and forepaw movement data were collected in *Fmr1^-/y^*(KO), EMX1-*Fmr1* KO, and WT mice (p30–100) in response to a grating stimulus at a fixed orientation presented at a variety of frequencies of phase-reversals separated by 10s gray screen intervals. **B.** LFP data was partitioned into “onset” periods (the first 250 msec following stimulus onset, stereotyped across stimuli) and “entrainment” periods (the remaining duration of the stimulus, varied based on the phase-reversal frequency). **C.** In both genotypes, N1 magnitude was highly correlated with the phase of the band-passed 2-10 Hz signal at the time of stimulus onset. **D.** In both KO and EMX1-*Fmr1* KO, the magnitude of the onset response was reduced relative to littermate WT mice (n = 37 p.g. for KO vs. WT littermate, n = 15 and 29, respectively, for EMX1-*Fmr1* KO vs. WT). Dots at bottom of plots in this figure indicate points of significant difference between groups, assessed by non-parametric hierarchical bootstrap with 99% confidence. **E.** There was no difference between genotypes in the amount of forepaw movement in the three seconds after stimulus onset. **F.** ssVEP amplitude (mean +/- SEM) for KO and WT littermate (left) and EMX1-*Fmr1* KO and WT (right) calculated for LFP data during the “entrainment” periods. * indicate temporal frequencies for which there is a significant difference between genotypes assessed through nonparametric hierarchical bootstrapping with a 99% confidence interval (CI). **G.** Bootstrapped difference (median +/- 99% CI) between genotypes in latency of the N1 of the VEP for each temporal frequency. Significant differences are found where the CI does not overlap with 0.

Pk1 oscillations (2-10 Hz) at the time of stimulus onset (Fig. 7C). We found the probability of a 180° phase occurring at stimulus onset was lower in *Fmr1^-/y^* mice (Extended Data 9A-B) and that the N1 amplitude appeared reduced across most phases in *Fmr1^-/y^* and EMX1-*Fmr1* KO mice (n = 15) (Fig. 7C, Extended Data 9C). Thus, the onset response was significantly smaller in *Fmr1^-/y^* and EMX1-*Fmr1* KO mice (Fig. 7D). There was no difference in the behavioral response to stimulus onset (Fig. 7E, Extended Data 9J).

We next analyzed the “entrainment” period in the frequency domain to calculate the steady-state VEP (ssVEP) amplitude (Extended Data 10A). ssVEP amplitude appeared enhanced within the range of periodic Pk1 for WT mice, and the amplitudes for *Fmr1^-/y^*and EMX1-*Fmr1* KO mice within this range were significantly smaller (Fig. 7F). Consistent with the “alpha” phenotype, evoked responses to stimuli phase-reversing at frequencies from 2 to 7.5 Hz were significantly delayed in *Fmr1^-/y^*and EMX1-*Fmr1* KO mice (Fig. 7G).

### Experience-dependent plasticity reduces genotype differences in VEP amplitude, but not latency, in a stimulus-specific manner

In a previous study, visually evoked theta oscillations in V1 of *Fmr1^-/y^*mice were shown to be altered only after induction of experience-dependent plasticity through repeated presentations of a grating stimulus^73^. Here, we have shown that alpha-like oscillations are altered in *Fmr1^-/y^* mice during resting-state measurements and that VEPs are reduced in amplitude relative to WT mice for unfamiliar visual stimuli. Perhaps impairments in experience-dependent plasticity in *Fmr1^-/y^* mice exacerbate genotype differences in VEPs. To test this, we selected one orientation at one phase-reversal frequency (either 7.5 Hz, n = 12 p.g., or 2 Hz, n = 17 p.g.) to present for either 2 days (for 7.5 Hz) or 6 days (for 2 Hz). On the final day, the now familiar oriented stimulus was interleaved with a grating of a novel orientation (Fig. 8A,E). We first measured behavioral habituation to the familiar stimulus and found robust orientation-selective habituation in *Fmr1^-/y^* mice for 7.5 Hz and 2 Hz stimuli (Fig. 8B,F)^74^. We separately analyzed onset and entrainment VEPs across days. For the 7.5 Hz stimulus, the reduced N1 of the onset response was replaced by an elevated positivity once the stimulus became familiar (significant difference 145-174 msec on days 2 and 3); this change was stimulus-specific (Fig. 8C). The positivity of the onset response to the familiar 2 Hz stimulus (days 2 through 7) was similarly elevated in *Fmr1^-/y^* mice (Fig. 8H), paralleling findings in FXS humans in the auditory domain^75^. Both genotypes showed enhanced “entrainment” responses to both the 2 and 7.5 Hz familiar stimuli, consistent with previously characterized stimulus-selective response plasticity (SRP)^59,74,76–78^, with no evidence of impaired plasticity in *Fmr1^-/y^* mice (Figure 8D,G). In fact, after the stimulus became familiar, there was no longer a significant genotype difference in the amplitude of the “entrainment” response (Fig 8C,K). However, latency differences between genotypes persisted across days: the N1 of the “entrainment” response to the familiar 2 Hz stimulus (days 2 through 7) was significantly increased in latency in *Fmr1^-/y^* mice, and the second negativity also appeared delayed on days 4 through 7 (Fig. 8J-K). SRP mechanisms are not impaired in *Fmr1^-/y^* mice; crucially, this motivates exploring therapeutics that might harness such plasticity mechanisms to correct electrophysiological alterations observed in FXS.

**Figure 8.**
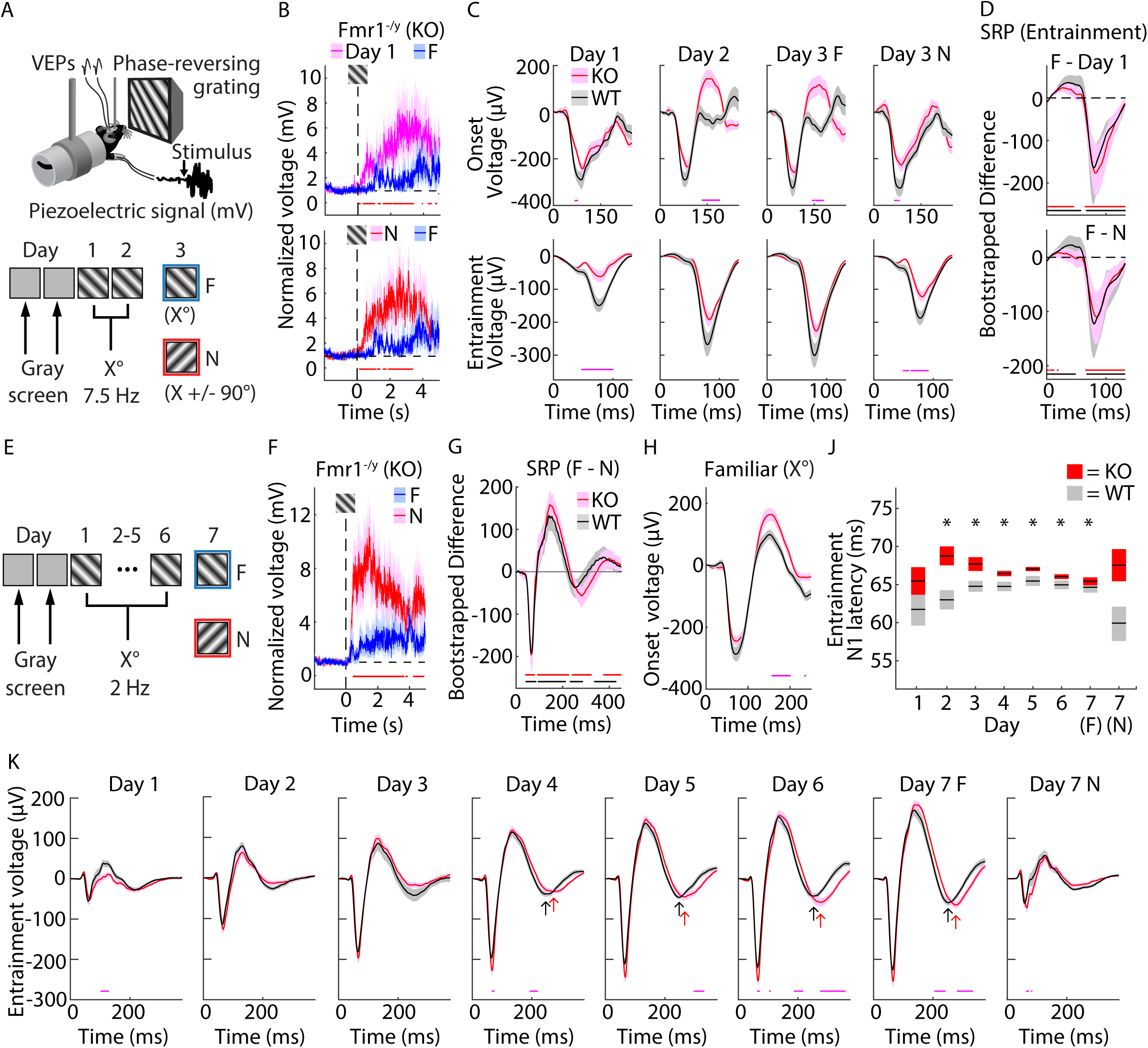
SRP reduces genotype differences in amplitude, but not latency, of VEPs. **A.** Experimental design. LFP data from L4 of V1 and forepaw movement data were collected in juvenile *Fmr1^-/y^* (KO) and WT mice (p30-40) in response to a grating stimulus phase-reversing at 7.5 Hz. After habituation to head-fixation for two days, the grating stimulus at a fixed orientation was presented for two days. On the third day, the familiar (F) orientation and a novel (N) orientation were presented in a randomly-interleaved fashion. **B.** Forepaw movement in response to stimulus onset in KO on days 1 and 3 (n = 12). Dots at bottom of plots in this figure indicate points of significant difference between groups, assessed by non-parametric hierarchical bootstrap with 99% confidence. **C.** Changes in the “onset” and “entrainment” period responses (mean+/- SEM) across days in KO and WT mice (n = 12 p.g.). **D.** Bootstrapped difference (median +/- 99% CI) between the “entrainment” responses for F and Day 1 (top) and F and N (bottom) for KO and WT, which are measures of stimulus-selective response potentiation (SRP). Red dots at bottom of the plots indicate points of significant difference for KO, black dots indicate these points for WT. **E.** A separate cohort of KO and WT mice (p30-100) were shown the same oriented, grating stimulus phase-reversing at 2 Hz for six days, then shown the F and N orientations on day 7. **F.** Forepaw movement in KO mice in response to onset of the F and N stimuli on day 7 (n = 17). **G.** Bootstrapped difference (median +/- 99% CI) between the “entrainment” responses to the 2 Hz F and N stimuli for KO and WT. Red dots at bottom of the plots indicate points of significant difference for KO, black dots indicate these points for WT. **H.** Onset responses to the familiar orientation, randomly sampled across days 2-7 for the familiar stimulus (n = 17 p.g.). **J.** VEP N1 latency during the “entrainment” period (mean +/- SEM) for KO and WT littermate mice across days. * indicate days for which there is a significant difference between genotypes assessed through nonparametric hierarchical bootstrapping with a 99% confidence interval (CI). **K.** Changes in the “entrainment” period VEP (mean+/- SEM) across days. A delayed second negativity in KO mice (red arrows) vs WT mice (black arrows) seems to appear days 4-7.

## Discussion

Disruptions in alpha oscillations are characteristic of neurodevelopmental and neuropsychiatric disorders, including FXS, wherein alpha oscillations are slowed. We have discovered a correlate for this FXS “alpha” phenotype in V1 of *Fmr1^-/y^* mice. Since frequency band analysis of absolute spectra can be misleading when studying conserved periodic signals across species, we isolated the periodic component of resting-state EEG data. This revealed that the shift in center frequency of periodic Pk1 in FXS is conserved in V1 of *Fmr1^-/y^*mice. Alpha oscillations are strongest over occipital cortex in humans, so conservation of the phenotype in V1 across species makes sense^51,52^. This discovery is an important step forward for at least two reasons: (1) it justifies the use of mice to dissect the cause(s) of alpha-band oscillatory EEG phenotypes observed in humans and gain better understanding of how cortical network activity goes awry in disease states, and (2) it begins to validate a putative transspecies biomarker that can be used to objectively quantify and compare putative treatment responses in mice and humans with FXS.

A key variable we found to influence the phenotype is age. We found that in children with FXS, Pk1 is both shifted to a lower frequency and reduced in maximum power relative to age-matched controls. The difference in center frequency is exaggerated in adults, but the maximum power is not significantly different from adult controls. These findings suggest that the age-dependent increase in Pk1 center frequency observed in typical development^48–50^ might stagnate in FXS, consistent with findings in ASD^40^. In V1 of *Fmr1^-/y^* mice viewing a static gray screen, we observed that the frequency shift within periodic Pk1 is also present and exaggerated in adults, and the Pk1 power differences are larger in juveniles. Unlike humans, the developmental change in Pk1 frequency appears to be from a decrease in center frequency in adult *Fmr1^-/y^* mice.

Nevertheless, the mouse appears to recapitulate different developmental windows of the alterations to Pk1. Since these alterations may have multiple mechanistic bases, efficacy of therapeutics could vary within different developmental windows^79^ and *Fmr1^-/y^*mice may be useful to optimize the timing of treatments. Future resting-state recordings in younger mice (p21) might reveal further developmental differences^29,80^.

In mice, another key variable that influenced the genotype differences in periodic Pk1 was the level of background luminance. We found that luminance amplified the genotype effect relative to when the mice were in the dark because it suppressed Pk1 power in *Fmr1^-/y^* mice, especially in juveniles, but enhanced Pk1 power in WT mice. In neurotypical humans, alpha oscillations are strongest in the dark or when eyes are closed^81,82^. This difference could arise because mice are nocturnal while humans are diurnal. Regardless, it will be of interest to test in future studies if alterations in Pk1 are amplified when FXS subjects close their eyes or are recorded in the dark.

The great utility of studying the mouse model of FXS is that it offers the opportunity to enter the brain to dissect the underlying basis for an electrophysiological phenotype. In the current study, for example, intracortical recordings revealed altered temporal dynamics and power differences that were not detectable at the surface. We note that intracortical recordings in *Fmr1-*KO rats have similarly shown reduced power of 3-9 Hz oscillations^27^. Our L4 recordings revealed a two subpeak structure of Pk1, with the frequency shift (“alpha” phenotype) in *Fmr1^-/y^* mice in the first subpeak (Pk1a). Previously, theta oscillations in WT mice have been analyzed as a single entity across this frequency range, giving rise to differing opinions as to which population of inhibitory interneurons coordinates these oscillations^51,54,83^. Our discovery of the two subpeak structure has offered some clarity: inhibition of SOM+ and PV+ neurons independently affects Pk1a and Pk1b, respectively. A mechanistic distinction in subpeaks might translate to humans, given reports of differential functionality of the lower and upper portions of the alpha frequency band^19,84,85^.

Our study is the first to experimentally relate hypoactivity of PV+ cells to the “gamma” phenotype of FXS. Elevated gamma power is also characteristic of other neurodevelopmental and neuropsychiatric conditions, including ASD and schizophrenia^21,39,86^. Inhibiting PV+ cells in WT mice reproduces both elevated periodic power seen *Fmr1^-/y^* mice and elevated aperiodic power seen in the humans with FXS, suggesting a uniform mechanistic basis across species. Presumably, the paradoxical increase in gamma power after PV+ suppression relates to disinhibition of reciprocally connected excitatory cells^87–90^.

However, hypoactivity of PV+ cells cannot fully account for the alterations in periodic Pk1. Instead, we find that inhibition of SOM+ interneurons phenocopies the alterations in Pk1a power observed in *Fmr1^-/y^* mice. Consistent with a hypothesis of diminished SOM+ cell activity in FXS, coupling of Pk1a oscillations to 14-38 Hz oscillations (regulated by SOM+ interneurons^58,67^) is weaker in *Fmr1^-/y^* mice. Further, induction of SRP, which recruits SOM+ interneurons^59^, corrects the reduced amplitude of VEPs in *Fmr1^-/y^* mice. Taken together, the data suggests that the FXS LFP phenotypes might reflect hypoactivity of both SOM+ and PV+ interneurons. Intriguingly, both classes of inhibitory interneurons are inhibited by vasoactive intestinal peptide-expressing neurons, which are documented as hyperactive in *Fmr1^-/y^* mice^91–94^.

Despite converging evidence that the alterations in periodic power in *Fmr1^-/y^* mice report differences in network inhibition, manipulating PV+ and SOM+ cells in WT mice fails to reproduce the shift in center frequency of Pk1a seen in *Fmr1^-/y^* mice. Similarly, induction of SRP only normalizes VEP amplitudes, not the increased latencies, in *Fmr1^-/y^* mice. This suggests the alterations in periodic Pk1 do not solely reflect changes in inhibition and might instead emerge from altered corticothalamic activity, given the role of the thalamocortical loop in alpha oscillations^34,52,65^.

In fact, it appears that the cause of all reported LFP phenotypes is the loss of FMRP in excitatory neurons and/or glia. This suggests that inhibition may be altered as a secondary consequence of changes in excitatory neuronal function^95,96^ and informs therapeutic approaches. For example, restoring FMRP expression only in cortical inhibitory interneurons might be misguided. Reexpressing FMRP in cortical excitatory neurons is sufficient to alter PV+ activity^97^. Future work will hopefully elucidate how loss of FMRP affects excitatory neuron activity *in vivo*, clarifying whether it drives hyperexcitability^5,98^ or hypoactivity^27,95^ of principal cells, and how this altered activity affects alpha-like oscillations.

Identifying the “alpha” phenotype in *Fmr1^-/y^* mice positions it as a candidate translational biomarker of FXS and offers the possibility of a parallel measure of drug efficacy across species to improve screening of therapeutics preclinically. Until recently, human alpha oscillations were considered a signal of cognition that could not be studied in mice, and now we have discovered that disruptions in alpha oscillations in FXS are conserved in alpha-like oscillations of *Fmr1^-/y^*mice. This fuels research into the underlying mechanisms of alpha oscillations and their disruptions to inform new therapeutic avenues, facilitates probing if therapeutics correct these disruptions, and increases confidence that successful correction of the disruptions preclinically will translate to success in humans in clinical trials. We hope our framework for studying alpha disruptions preclinically can be applied to research on other neuropsychiatric and neurodevelopmental disorders with pathological alpha oscillations to broadly advance translational research.

### Materials and Methods Human subjects

#### Participants for Pediatric FXS study

Resting-state EEG (rsEEG) data were collected from 17 males (27-78 months old) with full mutation of *FMR1* and 17 age-matched (27-80 months) typically-developing males. Participants were recruited as part of two studies (IRB-P00034676 and IRB-P00025493) conducted at Boston Children’s Hospital/Harvard Medical School. An additional secondary analysis IRB was approved for combining and analyzing data from both studies. FXS participants all had documented full mutation of the *FMR1* gene, however methylation status was not known for many participants, and participants were not excluded for size mosaicism (mixture of full and premutation). Additional exclusion criteria included history of prematurity (<35 weeks gestational age), low birth weight (<2000gms), known birth trauma, known genetic disorders (other than FXS), unstable seizure disorder, current use of anticonvulsant medication, and uncorrected hearing or vision problems.

#### Participants for Adult FXS study

rsEEG data were collected from 20 males (19-43 years old; M = 31.09, SD = 7.67), 20 with a full mutation of *FMR1* and 20 age-matched (20-44 years old; M = 29.42, SD = 7.12) typically-developed males. Participants were recruited as part of a study (IRB #2013-7327) conducted at Cincinnati Children’s Hospital and Medical Center. Adult FXS participants all had full FMR1 mutations (>200 CGG repeats), confirmed via past testing results made available in a participant’s medical record or via Southern Blot and/or PCR conducted in collaboration with the Molecular Diagnostic Laboratory at Rush University. Participants were not excluded for size mosaicism (mixture of full and premutation). Additional exclusion criteria for adult participants included known syndromic conditions associated with intellectual disability (other than FXS; e.g., Down Syndrome), being on benzodiazepines or anticonvulsant medications, or being on any novel potential treatment (e.g., minocycline) known to affect EEG measures within 4 weeks of testing. Typically developed controls were excluded if they scored ≥ 8 on the Social Communication Questionnaire (SCQ; Rutter, Bailey, & Lord, 2003), had a history of psychiatric or neurologic disorders, or had a first or second-degree relative with autism spectrum disorder or a serious psychiatric illness.

Institutional review board approval was obtained prior to starting each study. Written, informed consent was obtained from either the participant or from the participant’s parent/guardian with assent from the participant. For the pediatric study, written, informed consent was obtained from all parents or guardians prior to their children’s participation in the study.

We note that our study focused on male subjects (both humans and mice). The *Fmr1* gene is on the X-chromosome, FXS diagnoses are most prevalent in males, and the electrophysiological phenotypes are most prominent in the male subjects^19^. However, we recognize that there are sexually dimorphic phenotypes in FXS, including in alpha oscillations^18,55^, and encourage future study of the “alpha” phenotype in human females with FXS and in female *Fmr1^-/y^* mice.

### Human rsEEG data collection and preprocessing

#### Pediatric study

rsEEG data were collected while the child either sat in their caregiver’s lab or independently in a chair situated in a dimly lit, sound-attenuated, electrically shielded room. Continuous EEG was recorded while the participant was shown a silent screen saver of abstract colorful moving images for 2-5 minutes depending on the child’s compliance. In some cases, participants watched a silent video of their choosing to improve behavioral cooperation.

rsEEG data were collected using a 128-channel Hydrocel Geodesic Sensor Net (Version 1, EGI Inc, Eugene, OR) connected to a DC-coupled amplifier (Net Amps 300, EGI Inc, Eugene, OR). Data were sampled at 1000Hz and referenced to a single vertex electrode (Cz). Raw NetStation (NetStation version 4.5, EGI Inc, Eugene, OR) files were exported to MATLAB (version R2017a) for pre-processing using the Batch EEG Automated Processing Platform (BEAPP)^99^ with integrated Harvard Automated Preprocessing Pipeline for EEG (HAPPE)^100^. Preprocessing has previously been described in detail for similar data^101^. Briefly, data were 1 Hz high-pass and 100 Hz low-pass filtered, down-sampled to 250 Hz, and then run through the HAPPE module for 60 Hz line noise removal, bad channel rejection and artifact removal using combined wavelet-enhanced independent component analysis (ICA) and Multiple Artifact Rejection Algorithm (MARA) ^102^. Given the short length of EEG recording, 39 of the 128 channels were selected for ICA/MARA (Standard 10-20 electrodes: 22, 9, 33, 24, 11, 124, 122, 45, 36, 104, 108, 58, 52, 62, 92, 96, 70, 83; additional electrodes: 23, 28, 19, 4, 3, 117, 13, 112, 37, 55, 87, 41, 47, 46, 103, 98, 102, 75, 67, 77, 72, 71, 76). These electrodes were selected based on their spatial location, covering frontal, temporal, central, and posterior regions of interest (see Fig. 1). After artifact removal, channels removed during bad channel rejection were interpolated, data were rereferenced to an average reference, detrended using the signal mean, and segmented into 2-second segments. Any segments with retained artifact were rejected using HAPPE’s amplitude and joint probability criteria. EEG were rejected for data quality if they had fewer than 10 segments (20 seconds total) or did not meet the following HAPPE data quality output parameters: percent good channels >80%, mean and median retained artifact probability <0.3, percent of independent components rejected <80%, and percent variance after artifact removal <32%.

#### Adult study

rsEEG was continuously recorded and digitized at 1000 Hz, filtered 0.01-200 Hz, referenced to Cz, and amplified 10,000x using a 128-channel saline Electrical Geodesics system (EGI, MagStim, Minnesota) with sensors placed approximately according to the International 10/10 system. Data were preprocessed using MATLAB 2021b and the Cincinnati Very High Throughput Pipeline (VHTP) which utilized EEGLAB 2021b^103^.

rsEEG data used for the spectral power analysis were digitally filtered from 0.5 to 100 Hz with a 60 Hz notch (57 – 63; harmonics were removed up to the Nyquist frequency of the original sampling rate), channels with poor quality data were interpolated (no more than 5% of channels interpolated), segments of poor-quality data were manually rejected, and then remaining data were submitted to independent components analysis via EEGLAB for artifact removal. Artifacts (e.g., muscle tension, ocular-related events, heart rate, etc.) were removed as components and then data were average referenced and segmented into 2-second segments. During the creation of segments, data were submitted to an automatic amplitude rejection threshold where segments containing artifact exceeding +/-120 µV were automatically rejected.

For analysis of burst dynamics, data were also digitally filtered from 0.5 to 100 Hz with a 60 Hz notch (57 – 63; harmonics were removed up to the Nyquist frequency of the original sampling rate), with no more than 5% of channels interpolated for bad data, but data did not undergo manual segment rejection. Instead, a section of continuous data was isolated starting from approximately 60 seconds into the recording through approximately 230 seconds into the recording, depending on the degree of artifact present. The continuous segment was then submitted to ICA for artifact correction where removal of low frequency artifacts (e.g., heart rate, ocular-related events, etc.) was prioritized due to the planned low frequency filtering for burst analyses (i.e., high frequency noise was filtered out). Data were average referenced and then reduced to 90-second-long segments for analysis, as 90 seconds was approximately the longest segment of clean, continuous data available for analysis.

### Murine Subjects

*Fmr1^-/y^* mice were obtained from Jackson Laboratories, Maine, USA (stock # 003025) and backcrossed onto a C57BL/6J background for at least six generations at Massachusetts Institute of Technology or King’s College London. *Fmr1* cKO mice were also obtained from Jackson Labs (stock # 035184) and crossed with Emx1-Cre mice (Jackson stock # 005628). As previously reported, for hM4D(Gi) experiments, mice were Parvalbumin-Cre recombinase knock-in mice (B6;129P2-*Pvalb^tm1(cre)Arbr^*/J, PV-Cre) on a C57BL/6 background (Jackson stock # 017320)^66^. For optogenetic inhibition experiments, SOM-Cre mice were used (B6J.Cg-Ssttm2.1(cre)Zjh/MwarJ, Jackson stock # 028864). Experimental cohorts consisted of male littermates that were weaned at p21 and were p30-P150 at the time of experiments. Mice were maintained on a 12-hour light-dark cycle (7am – 7pm) with *ad lib* access to food and water and housed with 1-4 other littermates. All experiments were performed blind to genotype using age-matched littermate controls during the light phase. All experimental techniques were approved by The Institutional Animal Care and Use Committees at MIT or were in accordance with the UK Animals (scientific procedures) Act (1986), depending on the site where data were collected.

### Murine electrode implantation surgery

#### L4 LFP electrodes

Methods for LFP surgeries were followed as previously reported^59^. Briefly, mice (p30-150) were administered with pre-operative analgesia (0.1 mg/kg Buprenex subcutaneously) and anesthetized with isoflurane (3% in oxygen at induction, 1.5% in oxygen through operation). The head was shaved and disinfected with povidone–iodine (10% w/v) and ethanol (70% v/v), the scalp was incised, and the skull surface was scored. A steel screw for head-fixation was bound to the front of the skull with cyanoacrylate glue, the skull was levelled, and a reference electrode (silver wire, A-M systems) was placed in right frontal cortex. LFP tungsten recording electrodes (FHC), 75 μm in diameter at their widest point, were implanted in binocular visual cortex (-3.5 mm bregma, +/- 3 mm midline, 450 μm depth). All electrodes were secured using cyanoacrylate and the skull was covered in dental cement. A nonsteroidal antiinflammatory drug (meloxicam, 1.5 m/kg) was delivered subcutaneously post-operatively for 3 consecutive days and mice were monitored daily for discomfort. Mice were given 48-96 hours to recover before being habituated to head-fixation.

#### Cortical surface electrodes

Using similar preoperative and surgical techniques (except use of carprofen as the pre-operative analgesia), *Fmr1^-/y^* and littermate WT mice (P70-115) were chronically implanted with screw-type electrodes (1.6mm, Protech International) in the skull to measure cortical EEG in V1 and S1 and with the same type of LFP electrode as described above in V1. Screw electrodes were placed in burr holes over S1 (–2.06 mm bregma, 3.20 mm midline) and V1 (-3.50 mm bregma, –3 mm midline) with a reference electrode and a ground over the left and right olfactory bulb. The coordinate for the LFP electrode was the same as the skull screw (i.e., the same hemisphere) except the wire was inserted into the cortex tissue at a 10-degree angle and advanced in the dorsoventral axis to a depth of 0.52 mm from the cortex surface. Electrodes were connected to head-mounts and secured with dental cement and mice were given at least one week to recover before EEG/LFP recordings.

### Murine resting-state EEG/LFP data collection and preprocessing

Mice were habituated to the recording set-ups for at least two days. For habituation of head-fixed mice, the monitor displayed an iso-luminant gray screen (80 lux) generated using software written in either C++ for interaction with a VSG2/2 card (Cambridge Research Systems) or MATLAB (MathWorks) using the PsychToolbox extension (http://psychtoolbox.org). For resting-state LFP recordings in head-fixed mice, five minutes of data while the monitor was turned off were collected in some of the mice, and three to five minutes of data while the monitor displayed the iso-luminant gray screen were collected in all mice. Data were collected on a Plexon Recorder 64 system, with a PBX-211 2003 pre-amplifier. LFP data were collected from the electrode placed in V1 (left or right hemisphere) and forepaw movement data were collected from a piezoelectric disk placed under the animals’ forepaws; data in total were collected from three channels (the V1 channel, the ground in PFC, and the piezo). LFP voltage data were sampled at 1 kHz and band-passed through analog filtering via a low cut, 2-pole 3 Hz filter and a high-cut, 4-pole, 8 kHz filter. Frequencies down to 0.4 Hz were recovered using the inverse transfer function for the Linkwitz-Riley analog filter obtained from Plexon. This and other all other analyses were conducted using custom MATLAB code (R2018b). Data from the V1 LFP channel were loaded into MATLAB, unfiltered, converted to microvolts, and zero-meaned. A second-order IIR notch filter was applied to remove 60 Hz line noise with a bandwidth of 0.01 (for gray screen data) or 0.025 (for black screen data, wherein line noise was higher). Data from the piezo channel were also loaded into MATLAB and rectified.

For EEG recordings in freely-moving mice, the mice were tethered to four channel EEG/EMG recording systems (Pinnacle Technology Inc.) and housed individually and sequentially in a sound-proof and light-controlled cabinet equipped with a video camera. Voltage data were acquired continuously for a 10–12-hour period with a sampling rate of 1 kHz, amplified 100X, and band-passed filtered through analog filtering via a two-series, single pole simple RC filter at 1 Hz and an eight-order elliptical filter at 100 Hz. Frequencies down to 0.6 Hz were recovered using the inverse-transfer function for the passive RC filter obtained from Pinnacle and implemented in MATLAB (R2018b). 100-second-long resting state time series from S1 and V1 electrodes in each mouse were prepared using the synchronous video to identify when mice were in quiet wake (resting) vigilance state (i.e., sitting at the corner of the recording cage, moving the head but not the whole body, no running). This time-series data were loaded into MATLAB, unfiltered, converted to microvolts, and zero-meaned. A second-order IIR notch filter was applied to remove 50 Hz line noise with a bandwidth of 0.15.

### PV+ inactivation methods

As described in Kaplan et al. (2016) for hM4D(Gi) experiments, in addition to implanting LFP electrodes, V1 of p30-60 PV-Cre mice was infected with an AAV9-hSyn-DIO-HA-hM4D(Gi)-IRES-mCitrine virus (UNC viral core – generated by Dr. Brian Roth’s laboratory)^66^. Using a glass pipette and Nanoject system (Drummond scientific, Broomall, PA, US), 81 nL of virus was delivered at each of 3 cortical depths: 600, 450, and 300 µm from the cortical surface, with 5 min between re-positioning for depth. Mice were allowed 3–4 weeks recovery for virus expression to peak before experiments were initiated. After habituation, on the experiment day, five minutes of resting-state LFP data with mice head-fixed in front of a gray screen was collected before and 30 minutes after Clozapine-N-oxide (CNO, Enzo Life Sciences) was diluted in saline and injected intraperitoneally at a dosage of 5 mg/kg.

### SOM+ inactivation methods

For optogenetic inactivation experiments, we infected V1 of P40-50 SOM-Cre mice with either pAAV-Ef1a-DIO eNpHR 3.0-EYFP (AAV5) virus or the empty viral vector pAAV-Ef1a-DIO EYFP (AAV5). Using the aforementioned Nanoject system, we delivered 100 nL of virus at 2 nL/s at 4 cortical depths: 600, 450, 300 and 150 µm from the cortical surface, with 6 min between re-positioning for depth. In addition to implanting LFP electrodes and infusing the virus, ready-made optic fibers (200 µm girth) mounted in stainless steel optic cannulas (1.25 mm diameter, 2 mm fiber projection, Thor labs, CFMLC12L02, Newton, NJ, US) were then implanted lateral to the recording site and at a 22° angle to the recording electrode, 0.1 mm below surface in each hemisphere. 3 weeks later, mice were similarly habituated for two days.

For resting-state LFP data collection, while mice were head-fixed in front of a gray screen, we delivered five 10-s long continuous pulses of green light (550 nm, 6-7 mW) into V1 using a digitally-controlled LED driver (PlexBright® Optogenetic Stimulation System). Each pulse was separated by 60 sec, resulting in a recording session that lasted about 7 minutes.

### Cross-species resting-state EEG/LFP analysis Spectral analysis

Spectral analyses were performed in MATLAB (R2018b) using the Chronux toolbox^104^. For all human subjects, the power spectral density at each electrode, for each 2 second segment, was calculated with a multi-taper spectra analysis, using three orthogonal tapers and a time-bandwidth product (*TW*) of 2. The spectra were averaged across segments and then averaged across electrodes. The following electrodes were averaged for each region of interest (ROI): frontal - 3, 4, 11, 19, 23, 24, 28, 117, 124; central - 13, 36, 37, 55, 87, 104, 112; temporal - 41, 45, 47, 46, 52, 58, 96, 98, 102, 103, 108; posterior - 67, 70, 71, 72, 75, 76, 77, 83. All listed electrodes from each ROI were averaged for the full brain absolute power spectrum.

For freely-moving murine subjects, the 100 sec resting-state time series was segmented into 5 sec segments, resulting in 20 segments per subject. Longer segments were needed for the mice as the periodic signals of interest were at lower frequencies. The appropriate time-bandwidth product was calculated (*TW =* 2\**FR* /segment length)^105^ to achieve a consistent frequency resolution (*FR*) with human data (2 Hz). Multi-taper spectra were calculated with 9 tapers and a time-bandwidth product of 5, with no zero padding. For head-fixed murine subjects with L4 LFP electrodes, the last 150 sec of resting-state and piezo data while the monitor was off or displaying a static gray screen were extracted for spectral analysis. 5 sec segments were again used, but due to occasional contamination of the piezo signal into the LFP signal during certain movement bouts, a very small fraction of segments (usually 0-2 per mouse) were eliminated if they contained both a large movement artifact and a huge spike in low frequency power (less than 1.5 Hz). As a result, 28-30 segments per animal per condition were usually analyzed. The multi-taper spectra were calculated using 9 tapers and a time-bandwidth product of 5, as before. For additional analysis of 2-10 Hz data, multi-taper spectra were also calculated with a higher frequency resolution (1 Hz) by using 4 tapers and a time-bandwidth product of 2.5.

### Modified one over F fitting analysis

An effective way to model the 1/f signal of the absolute power spectrum across a variety of datasets is to use a Lorentzian function^47^. Published code (SpecParam, https://fooof-tools.github.io/fooof/) allows users to fit a Lorentzian to a spectrum, optimized for selection of a fitting range with no periodic power at the edges of the range^47^. Realistically, selecting such a fitting range is challenging with biological data, and indeed, the implementation of the Lorentzian fit through SpecParam does not work well for pediatric rsEEG data^48^. The algorithm cut through the absolute spectrum of the pediatric data in the middle of the fitting range, generating negative periodic power between 10-20 Hz, which is not theoretically possible. This issue was resolved previously by shifting the fit downwards (changing the y-intercept but not the slope) until there was no more negative periodic power^48^. In the present study, we encountered multiple issues of overestimated aperiodic power when using SpecParam to fit the 1/f component of both human EEG data of all ages and mouse LFP data, especially at the bottom edge of the fitting ranges and in the 10-20 Hz range (Extended Data 1B, Extended Data 4B). We ran SpecParam in MATLAB R2018b using the MATLAB wrapper (https://github.com/fooof-tools/fooof_mat) and used the following settings: peak width = [0.5 18], number of peaks = 7, peak threshold = 1, and ‘fixed’ mode for EEG data and ‘knee’ mode for LFP data. Since our L4 LFP data in mice did not have a linear aperiodic component (Extended Data 4A-B), the fits could not be uniformly adjusted in the same way as before.

Therefore, we wrote custom code in MATLAB (R2018b) to fit Lorentzian functions to the 1/f signals effectively and consistently across all our datasets. Our implementation required the user to have confidence that only one, not both edges of their selected fitting range be without periodic power, which is more realistic given limitations of experimental datasets (such as artifacts, filters, etc.). The first step, as outlined previously, was to plot the average power spectra per group in log-log space^47^. This allowed us to identify a good fitting range and to decide if the 1/f fit will be linear or nonlinear (which affected the fitting procedure but maintained consistent application of math and methodology). Our fitting is performed in log-log space on the average absolute spectrum for each subject. Our approach is more data-driven than SpecParam, but this minimized negative periodic power in a consistent way across all datasets.

For data requiring linear fits in log-log space, which was all data recorded outside of cortex (human and mouse EEG), the bottom edge of the fitting range was determined by the lowest frequency unaffected by high pass filtering (2 Hz for humans, 3 Hz for mice). The [x,y] coordinate for the first point of the fit was [edge of fitting range, average power of the lowest 0.6 Hz of the fitting range]. Next, the algorithm identified the frequency value above 8 Hz that yielded the largest slope magnitude for the fit, as shallower slopes would cut through the absolute spectrum and generate negative periodic power. The slope was calculated using this frequency value (x) and the average power at that frequency and at the frequencies on either side of it (y) as the second coordinate. The Lorentzian was then subtracted from the data, revealing the periodic spectrum. To analyze the periodic peaks for human data, we found the maximum values of the periodic spectra less than (Pk1) and greater than (Pk2) 15 Hz. The x coordinate of these maximum values was the center frequency, and the y coordinate was the maximum power of the peak. For mouse EEG data, there was only one periodic peak, which was identified as the maximum value below 10 Hz through a locally-weighted smoothing linear regression filter.

For data with a non-liner 1/f signal in log-log space, which was all the L4 LFP electrode data, an additional “knee” parameter was needed in the Lorentzian function. Our general approach for fitting the “knee” was consistent with SpecParam, where we first fit a linear aperiodic component within the frequency range above the knee and then fit the knee to the low-frequency values below the inflection point^47^. Thus, we needed to fit our initial linear 1/f estimate based on the upper edge of the frequency range, which meant it was critical to select an upper point with no periodic power. This required fitting out to 175 Hz, as there was still periodic power (especially in *Fmr1^-/y^* mice) through about 160 Hz. The frequency value (x_1_) with the lowest absolute power (ignoring the 45-75 Hz range if there was notch filtering) and the average power of it and its neighboring two frequency values (y_1_) were used to form the first coordinate for the linear aperiodic estimate. Then, the algorithm identified the frequency value between 9 and 25 Hz (one of the problematic ranges for SpecParam) that yielded the smallest magnitude slope of the linear fit, as steeper slope values would cut through the absolute spectrum and generate negative power. The second coordinate for linear estimate was determined from this frequency value (x_2_) and the average power at that frequency and at the frequencies on either side of it (y_2_). The slope and intercept values of the Lorentzian were ascertained through this linear estimate. To determine the knee, we found the local minima power values at frequencies below 2.24 Hz (below the first periodic peak), and, assuming these values were less than the linear 1/f estimate at this frequency, used them to calculate multiple knee value estimates according to the Lorentzian function: (10^(b-y))-(x.^(-1*m)) where (x,y) are the (frequency, power) coordinates of the spectrum, m is the slope, and b is the intercept (offset). The final knee parameter was calculated as the average of these estimates, and with the full equation for the non-linear 1/f, we then subtracted it from the absolute spectrum to analyze the periodic spectrum. To analyze the subpeaks of periodic Pk1 (2-10 Hz), we found the largest local minimum of the data in the middle of this range using a locally-weighted smoothing linear regression filter. Pk1a was either the maximum value of the periodic spectrum below the local minimum or the maximum value below 6 Hz, and Pk1b was the maximum value above the local minimum or 6 Hz.

### Phase-amplitude coupling analysis

Following previously published methods, modulation index was calculated from the probability distribution of the amplitude signal across 18 bins of phases^54,106^. Signals were band-passed using the *eegfilt* function in MATLAB designed by Scott Makeig (https://sccn.ucsd.edu/~arno/eeglab/auto/eegfilt.html) and then the Hilbert transform was calculated to find the amplitude or phase values. To generate comodulagrams, the phase signal was analyzed from 2 to 7 Hz with a step size of one and a bandwidth of two, while the amplitude signal was analyzed from 12 to 60 Hz with a step size of two and a bandwidth of four.

### Burst analysis

#### Murine subjects

To analyze the temporal dynamics of periodic Pk1, the same bandpass filter and Hilbert transform methodology was used, filtering 100 sec of continuous LFP or EEG data between 2-10 Hz. If more than 100 sec of continuous data were available, the starting point of the 100 sec chunk was randomly generated. Following published methods, a burst threshold was set at the 90^th^ percentile of the bandpass amplitude for each animal^55^. Burst count was quantified as the number of threshold crossings, and burst duration was the length of time spent above the threshold in one crossing; a threshold crossing had to last for more than 3 msec to count as a burst^55^. For animals with piezo data, movement bouts were identified where the piezoelectric signal exceeded a voltage of 2mV (standard threshold across all animals). To compensate for voltage drops below 2mV in the middle of movement bouts, a buffer threshold of 750ms was used. The voltage signal was subsequently converted to binary movement state categorization (1 = moving, 0 = still) and aligned to the band-passed amplitude signal to determine if bursts occurred during intervals of movement or quiescence. The sum of this binary signal/(100 sec*sampling frequency) was used to determine total % time spent moving in a session.

#### Human subjects

Burst counts and lengths were calculated from the same selection of occipital channels used for the spectral power analysis, but using the 90 s of continuous (non-segmented) data. Each individual channel was filtered according to a frequency range determined in a data-driven manner from the analysis of Pk1 in the periodic spectrum. Given the shift in center frequency of Pk1 for FXS subjects, data were filtered from 5-10 Hz for FXS and 7-12 Hz for TDC. Filtered data were Hilbert transformed and then converted to the absolute value of the log-transformed squared magnitude (i.e., power). As with the mice, a 90^th^ percentile threshold for each individual was used to calculate endogenous burst counts and burst lengths for each channel. Individual channel burst metrics were averaged across channels to provide the average burst dynamics over the occipital region of the head.

### Murine visual stimulus delivery

All visual stimuli, including the iso-luminant gray screen described above, were generated using previously published code (https://github.com/jeffgavornik/VEPStimulusSuite). After collecting gray screen resting-state LFP data, a subset of head-fixed mice (p30-100) were shown an oriented phase-reversing grating stimuli that ramped through seven temporal frequencies. These visual stimuli consisted of full-field, 100% contrast, 0.5 cycles per degree, sinusoidal gratings that were presented on a computer monitor. Grating stimuli spanned the full range of monitor display values between black and white, with γ-correction to ensure gray-screen and patterned stimulus conditions were iso-luminant. Each temporal frequency was presented at the same orientation with 50 phase reversals, and different frequencies were separated by 10 s of gray screen. The ramp always started with the lowest temporal frequency (2 Hz) and ended with the highest (15 Hz), and the ramp repeated nine times total, producing a recording session of about 23 min of LFP data.

After seeing the temporal frequency ramp, a subset of the mice (p30-40) were shown a grating stimulus at a new orientation that phase-reversed at 7.5 Hz, presented in 5 blocks of 100 phase reversals separated by 30 sec of gray screen. The mice were first head-fixed in front of a static gray screen for 5 min before being shown the phase-reversing stimulus, resulting in an approximately 8 min long LFP recording session. Mice were shown this same stimulus again the next day, and on the third day, they were shown the familiar orientation interleaved with a novel one offset by 90°. A separate cohort of mice (p30-150) were shown a grating stimulus at a new orientation that phase-reversed at 2 Hz, also presented in 5 blocks of 100 phase reversals separated by 30 sec of gray screen. This LFP recording session lasted around 15 min and was repeated for six consecutive days. On the seventh day, the mice were shown the familiar orientation interleaved with a novel one offset by 90°.

### Visual-evoked data analysis

VEP analysis was adapted from published methods^59^. Data were separated into “onset” (initial 250 msec of a block of oriented grating stimuli) and “entrainment” periods (rest of the block). The magnitude of the first negativity (N1) after onset was calculated as the maximum negative value in the first 125 msec after stimulus onset. The instantaneous phase of the periodic Pk1 oscillations at the time of stimulus onset was calculated from the Hilbert Transform of the band-passed 2-10 Hz signal and then binned (18 bins). If the piezo voltage value crossed the movement threshold (2 mV, see above) during the 250 msec period, the onset was classified as “moving” instead of “still.” For the average VEPs, data was smoothed with a Gaussian spanning 20 msec using MATLAB’s *smoothdata* function. The latency to the N1 of the VEPs during the “entrainment” period was calculated as the time to the maximum negativity within the first 99 msec after each phase reversal. VEP amplitude was calculated as the absolute value of the N1 voltage + the maximum positive voltage value between 99 and 130 msec. For the temporal frequency ramp, ssVEPs were calculated by taking the Fourier transform of the signal (which was divided into 2 sec segments for 2, 4, 6, and 7.5 Hz stimuli, and 1 sec stimuli for 10, 12, and 15 Hz stimuli) and finding the amplitude and phase at the stimulation frequency for each segment^107^. Normalized ssVEPs were calculated by dividing the Fourier transform of the signal by the Fourier transform of equivalent (segmented) lengths of gray screen data. For visual-evoked movement data, 3 sec of piezo voltage signal following stimulus onset was normalized to 1.5 sec of movement data prior to stimulus onset. For measurement of orientation-selective response habituation, 5 sec of piezo voltage signal following stimulus onset was normalized to 2 sec of movement data prior to stimulus onset^74^.

### Statistics

Statistics were conducted in MATLAB (R2018b). Initial statistical tests using the one-sample, two-sided Kolmogorov-Smirnov test (MATLAB function *kstest*) confirmed non-normal distributions of the various datasets. Thus, statistical tests were conducted using methods that did not assume normality, including the two-sample, two-sided Wilcoxon Rank-Sum test (also known as the Mann-Whitney U-test, MATLAB function *ranksum*) and non-parametric hierarchical bootstrapping^59,108^. The former was used to test for group differences for metrics that could be well-summarized by a single (or average) value per animal/subject (i.e., aperiodic parameters, periodic peak parameters, microburst count, etc.), while the latter was used for more complicated datasets where a single-value summary failed to capture the variability in the data (i.e., time-series, VEP waveforms and their latencies/amplitudes, etc.). The output of the MATLAB *ranksum* function yielded a rank-sum statistic, a z-statistic for larger sample sizes, and a p-value. The effect size was determined by finding the absolute value of the z-statistic divided by the square root of the summed sample sizes, yielding a value between 0 and 1 (where values >= 0.3 indicated a medium to large effect size). This non-parametric test does not have degrees of freedom, and confidence intervals were assessed through non-parametric bootstrapping. Code for bootstrapping was written following published methods^59,108^. For bootstrapping, to maintain littermate pair comparisons, the random selection of trials and animals was matched between the distribution of WT and *Fmr1^-/y^* mice when their sample sizes matched. 95% confidence intervals were used for human data and 99% percent confidence intervals for mouse data when bootstrapping. Finally, for measurement of the linear correlations in human EEG data, Pearson’s r^2^ (which also does not assume normality) was determined by fitting a line to the data (MATLAB function *polyfit*) and then dividing the sum of the squared residuals by the the total sum of squares (i.e., the variance times the sample size minus one) and subtracting that value from 1.

The effect size (Pearson’s r) was determined by taking the square root of r^2^, yielding a value between 0 and 1 where values >= 0.5 were considered a large effect size and values <=0.1 were considered a small effect size.

### Data and Code Availability

All preprocessed resting-state data from mice and corresponding analysis code will be made publicly available on Figshare and Github, respectively, upon publication. All remaining data from mice will be readily available upon reasonable request from the authors.

For data from human subjects, consents obtained from human participants prohibit sharing of deidentified individual data without data use agreements in place. The current study includes data collected from several independent projects, including some in which consent for open sharing was not collected from the subjects. Thus, our ability to share data from human subjects depends on a variety of factors: which specific data are requested; whether and to what extent the participants included in the requested data have consented to the sharing and future use of their data; whether deidentified, limited, or identified data are requested; and the purpose for which the data are requested. Prior to sharing data, we may need to confirm whether Cincinnati Children’s Hospital and Medical Center or Boston Children’s Hospital and the institution of the individual requesting the data have existing agreements or subcontracts with terms of data use and sharing that define the collaboration (e.g., data use agreements, data access agreements, subawards, reliance agreements). Please contact the corresponding author with reasonable data requests to determine availability of the specific data of interest.

## Acknowledgements

The authors thank R. Komorowski, M. Fai-Fong, and A. Heynen for their contributions to early experiments in *Fmr1* KO mice and D. Stoppel for his expertise in and management of various *Fmr1* KO mouse lines. The authors also thank L. DeStefano for feedback on the project, J. Gavornik for creating the visual stimulus delivery code, and A. Palisano for administrative support. This work was supported by the following grants: NIH 1K23DC017983-01A1 (C.L.W), NIH 1T32MH112510 (C.L.W.), U54HD082008 (C.A.E), U54HD104461 (C.A.E), NIH K23 HD101416 (L.M.S), NIH F31 HD113221-01A1 (J.E.N.), NIH R01EY023037 (M.F.B.), NIH R21NS123499 (M.F.B.), NSF GRFP (S.S.K-S.), FRAXA Foundation (S.S.K-S. and C.L.W), Autism Science Foundation (C.L.W.), The Pierce Family Fragile X Foundation (C.L.W.), Thrasher Research Fund (C.L.W.), Harvard Catalyst Medical Research Investigator Training Award (C.L.W.), Simons Foundation SFARI 575135 (M.F.B.), the Picower Institute Innovation Fund (M.F.B), Wellcome (207727/Z/17/Z)(S.F.C) and the Biotechnology and Biological Sciences Research Council (BBSRC) (BB/S008276/1)(S.F.C).

## Contributions

S.S.K-S. and M.F.B. conceived the project, secured funding with P.S.B.F., and designed the experiments with input from P.S.B.F. and M.J.H. C.L.W. secured funding and collected and preprocessed all pediatric EEG data with assistance from M.K. and helped conceive cross-species analyses. L.M.S. and C.A.E. secured funding and collected all adult EEG data, which J.E.N. preprocessed in multiple ways and analyzed for burst dynamics with input from L.E.E. S.F.C. secured funding and C.G. collected all freely moving mouse EEG data with input from S.F.C. F.A.C. collected all optogenetics LFP data with input from S.F.C. P.S.B.F. and M.L. collected some head-fixed mouse data. S.S.K-S. collected most of the head-fixed resting state and visually evoked LFP data, conceived and wrote the MATLAB code for cross-species spectral analyses, burst analyses, and VEP analyses, and analyzed mouse and human data (after it was preprocessed). S.S.K-S. interpreted the data and wrote the manuscript with review and editing from M.F.B, C.L.W., C.A.E., L.E.E., J.E.N., C.G., P.S.B.F, L.M.S., and S.F.C.

## Ethics Declaration

The authors declare no competing interests.

