## Supplemental figures and legends for "A human electrophysiological biomarker of Fragile X Syndrome is shared in V1 of *Fmr1* KO mice and caused by loss of FMRP in cortical excitatory neurons"

**Extended Data 1 (corresponds to figure 1). 1/f fitting methodology and detailed cross-sectional developmental trajectories of periodic Pk1 and Pk2. A.** Average power spectra for rsEEG of both FXS and TD children and adults are best fit by a line in log-log space (linear 1/f). **B.** Methodology. Comparison of the fit generated with the SpecParam code[47] and our improved 1/f fit for power spectra from a representative subset of children and adult subjects, plotted in log-log space. Red arrows identify regions where the SpecParam fit overestimates the aperiodic power, which occurred consistently in the 3-6 Hz and 10-20 Hz ranges. Our data-driven approach fits the line to these problematic regions for an improved fit. **C.** Identification of the maximum power and center frequency for each TD (left) and FXS (right) child for periodic Pk1 (green) and Pk2 (blue). **D.** Distribution of Pk1 maximum power and center frequency for FXS (red) and TD children (black). **E.** Pk2 center frequency values as a function of age for the FXS children, demonstrating that the three outlier frequency values were measured in older subjects. **F-G.** Maximum power as a function of age for Pk2 (D) and Pk1 (E) for FXS and TD children with corresponding lines of best fit and $r^2$ values. For (D), effect size (Pearson's r) = 0.455 (FXS) and 0.021 (TD). For (E), effect size = 0.18 (FXS) and 0.307 (TD). **H-J.** Same as (C-D) but for FXS and TD adults. **K-L.** Boxplot (median, IQR, and full range) and individual data points for the center frequencies (H) and maximum powers (J) of Pk2 for FXS and TD adults, WRST z-statistic = 0.611, effect size = 0.097 for (H), WRST z-statistic = 0.69, effect size = 0.109 for (J). **M.** Line of best fit and linear regression $r^2$ value for the correlation between the maximum power of Pk1 and Pk2 for FXS (effect size = 0.359) and TD children (effect size = 0.5).

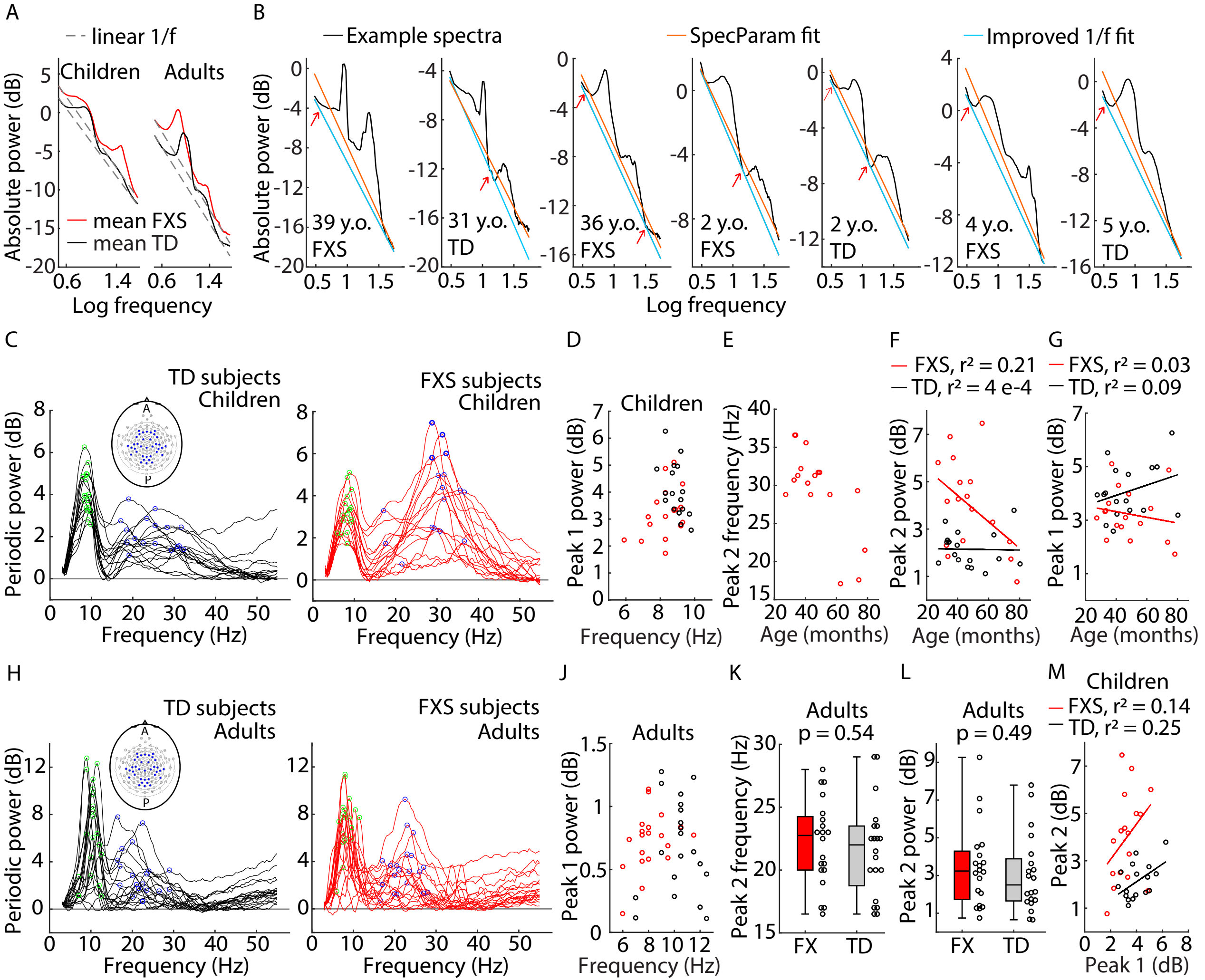

A
linear 1/f
Children
Adults
mean FXS
mean TD
Absolute power (dB)
Log frequency
B
Example spectra
SpecParam fit
Improved 1/f fit
39 y.o. FXS
31 y.o. TD
36 y.o. FXS
2 y.o. FXS
2 y.o. TD
4 y.o. FXS
5 y.o. TD
C
TD subjects Children
FXS subjects Children
Periodic power (dB)
Frequency (Hz)
A
P
D
Children
Peak 1 power (dB)
E
Peak 2 frequency (Hz)
Age (months)
F
FXS, r² = 0.21
TD, r² = 4 e-4
Peak 2 power (dB)
G
FXS, r² = 0.03
TD, r² = 0.09
H
TD subjects Adults
FXS subjects Adults
J
Adults
K
Adults
p = 0.54
FX
TD
L
Adults
p = 0.49
Peak 2 power (dB)
M
Children
FXS, r² = 0.14
TD, r² = 0.25
Peak 2 (dB)
Peak 1 (dB)

**Extended Data 2 (corresponds to figure 1A-K). rsEEG phenotypes of FXS children by ROI.** **A.** Aperiodic fit (mean +/- SEM) for electrodes in the frontal ROI for FXS and TD children. Dots at bottom of plots in this figure indicate points of significant difference between groups, assessed by non-parametric hierarchical bootstrap with 95% confidence. **B.** Boxplot (median, IQR, and full range) and individual data points for aperiodic offset values (the power at 3 Hz) from the fits in (A), WRST z-statistic = 2.239, effect size = 0.384. **C.** Periodic power spectrum (mean +/- SEM) for frontal electrodes. **D-E.** Boxplot and individual data points for the center frequency (D) and maximum power (E) of the Pk1 in the periodic spectrum in (C), WRST z-statistic = -1.971, effect size = 0.338 for frequency and WRST z-statistic = -1.412, effect size = 0.242 for power. **F-U.** Same as (A-E) but for electrodes in the central (F-K), temporal (L-P) and occipital (Q-U) ROIs. For (G), WRST z-statistic = 2.549, effect size = 0.437. For (J), WRST z-statistic = -1.942, effect size = 0.333. For (K), WRST z-statistic = -2.411, effect size = 0.414. For (M), WRST z-statistic = 2.239, effect size = 0.384. For (O), WRST z-statistic = -2.949, effect size = 0.506. For (P), WRST z-statistic = -1.894, effect size = 0.325. For (R), WRST z-statistic = 2.136, effect size = 0.366. For (T), WRST z-statistic = -1.824, effect size = 0.313. For (U), WRST z-statistic = -2.79, effect size = 0.479.

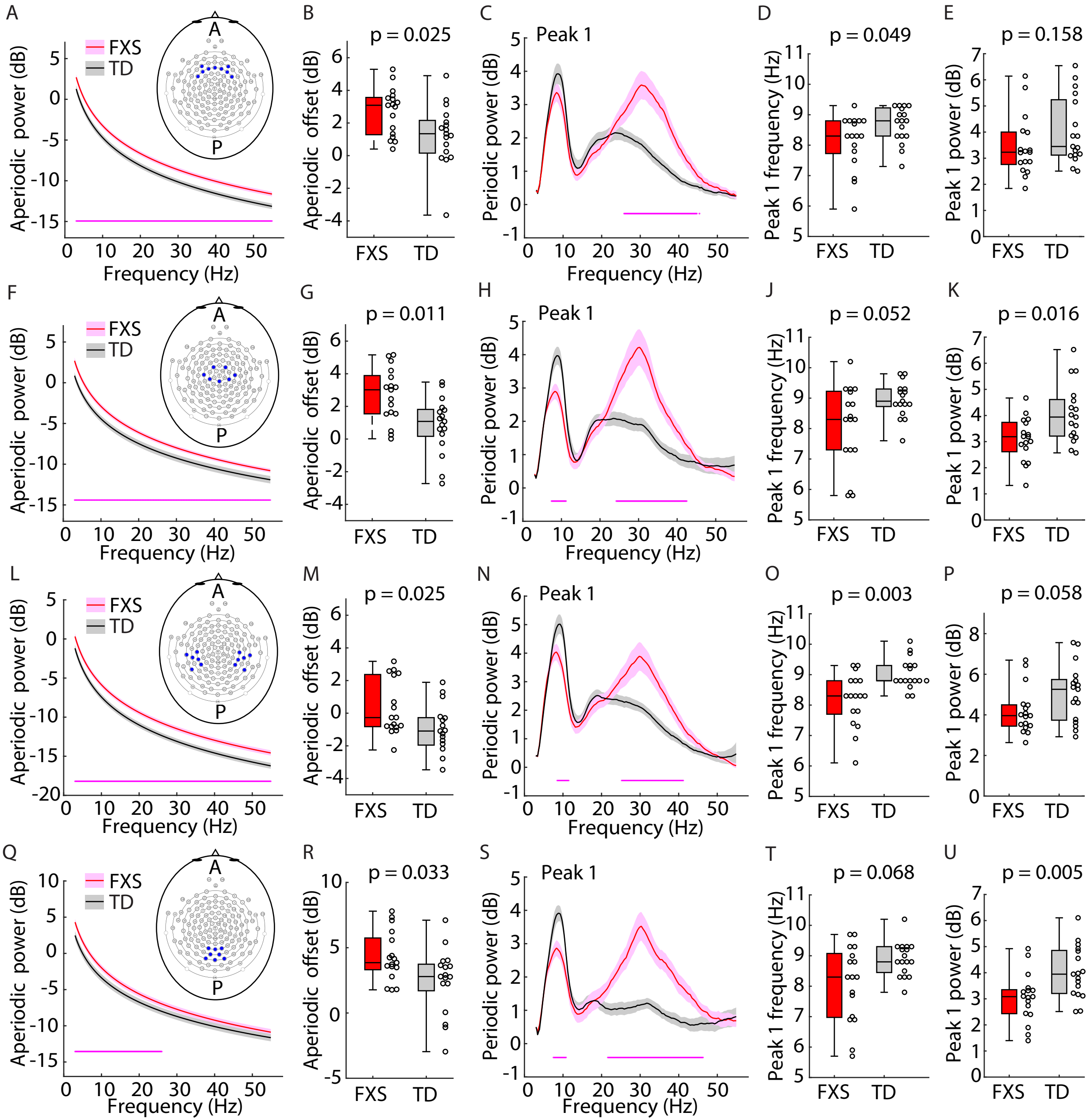

A
FXS
TD
A
P
Aperiodic power (dB)
Frequency (Hz)
B
p = 0.025
Aperiodic offset (dB)
FXS
TD
C
Peak 1
Periodic power (dB)
Frequency (Hz)
D
p = 0.049
Peak 1 frequency (Hz)
FXS
TD
E
p = 0.158
Peak 1 power (dB)
FXS
TD
F
FXS
TD
A
P
Aperiodic power (dB)
Frequency (Hz)
G
p = 0.011
Aperiodic offset (dB)
FXS
TD
H
Peak 1
Periodic power (dB)
Frequency (Hz)
J
p = 0.052
Peak 1 frequency (Hz)
FXS
TD
K
p = 0.016
Peak 1 power (dB)
FXS
TD
L
FXS
TD
A
P
Aperiodic power (dB)
Frequency (Hz)
M
p = 0.025
Aperiodic offset (dB)
FXS
TD
N
Peak 1
Periodic power (dB)
Frequency (Hz)
O
p = 0.003
Peak 1 frequency (Hz)
FXS
TD
P
p = 0.058
Peak 1 power (dB)
FXS
TD
Q
FXS
TD
A
P
Aperiodic power (dB)
Frequency (Hz)
R
p = 0.033
Aperiodic offset (dB)
FXS
TD
S
Peak 1
Periodic power (dB)
Frequency (Hz)
T
p = 0.068
Peak 1 frequency (Hz)
FXS
TD
U
p = 0.005
Peak 1 power (dB)
FXS
TD

**Extended Data 3 (corresponds to figure 1L-T). rsEEG phenotypes of FXS adults by ROI.** Same as extended data 2 but for FXS and TD adults. For (B), WRST z-statistic = 2.232, effect size = 0.353. For (D), WRST z-statistic = -3.89, effect size = 0.651. For (E), WRST z-statistic = 0.609, effect size = 0.096. For (G), WRST z-statistic = 2.232, effect size = 0.353. For (J), WRST z-statistic = -3.471, effect size = 0.549. For (K), WRST z-statistic = 0.23, effect size = 0.036. For (M), WRST z-statistic = 2.245, effect size = 0.387. For (O), WRST z-statistic = -3.341, effect size = 0.528. For (P), WRST z-statistic = 0.582, effect size = 0.092. For (R), WRST z-statistic = 2.367, effect size = 0.374. For (T), WRST z-statistic = -3.541, effect size = 0.56. For (U), WRST z-statistic = 0.392, effect size = 0.062.

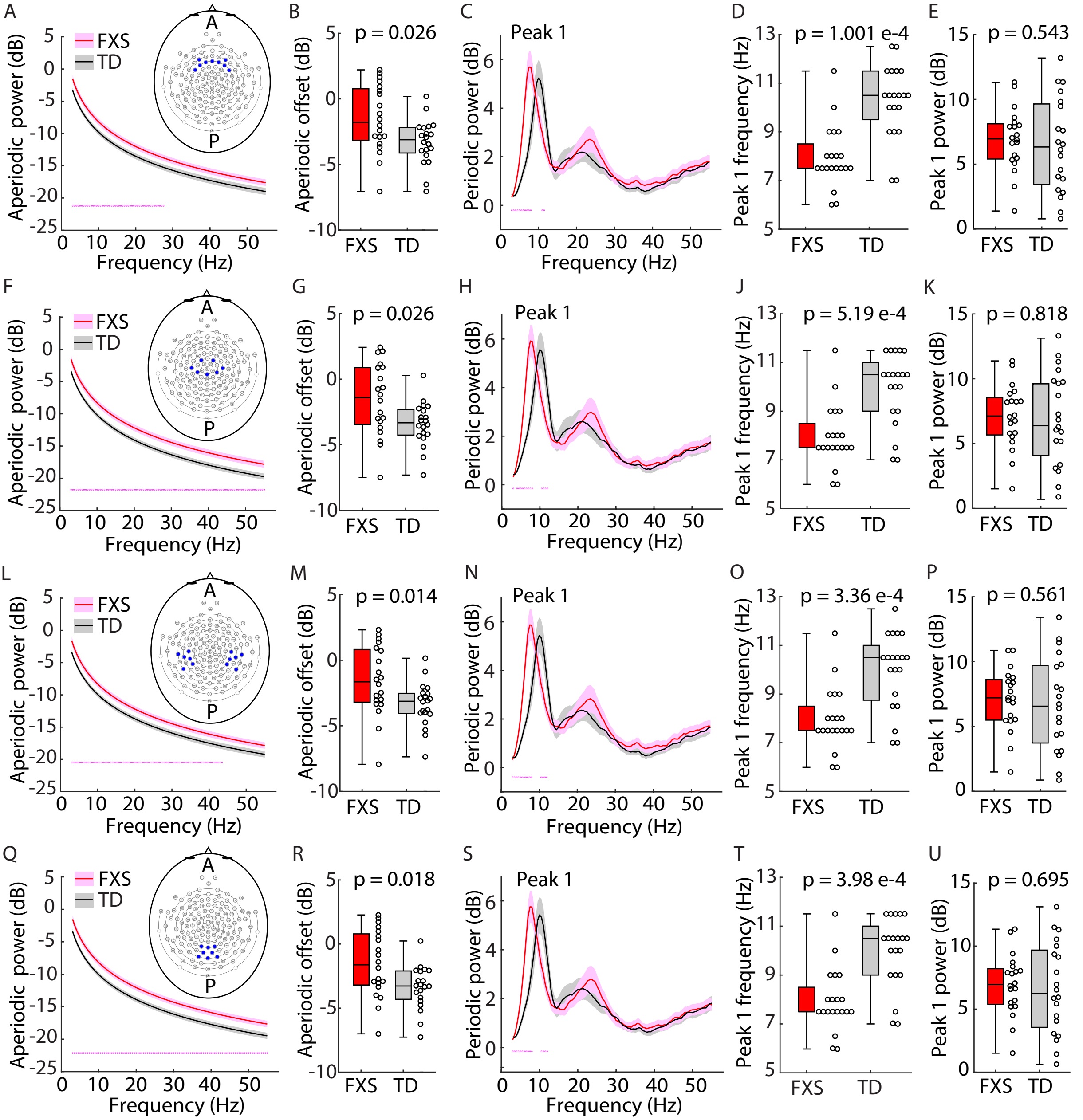
A
FXS
TD
Aperiodic power (dB)
Frequency (Hz)
A
P
B
p = 0.026
Aperiodic offset (dB)
FXS TD
C
Peak 1
Periodic power (dB)
Frequency (Hz)
D
p = 1.001 e-4
Peak 1 frequency (Hz)
FXS TD
E
p = 0.543
Peak 1 power (dB)
FXS TD
F
G
p = 0.026
H
J
p = 5.19 e-4
K
p = 0.818
L
M
p = 0.014
N
O
p = 3.36 e-4
P
p = 0.561
Q
R
p = 0.018
S
T
p = 3.98 e-4
U
p = 0.695

**Extended Data 4 (corresponds to figure 3). Aperiodic fits of L4 LFP data in V1 of mice and age-dependent phenotypes in Pk1 of *Fmr1$^{-/y}$* mice. A.** Methodology. Same as Fig. 3A but plotted in log-log space, revealing the nonlinearity of the 1/f. The inflection point of the curve is captured in the knee parameter. **B.** Comparison of the fit generated with the SpecParam code and our improved 1/f fit for power spectra from an example *Fmr1$^{-/y}$* (KO) and WT mouse viewing a black screen, plotted in log-log space. Red arrows identify regions where the SpecParam fit overestimates the aperiodic power, which occurred consistently in the 1.5-4 Hz and 10-20 Hz ranges, similar to the EEG data. Our data-driven approach fits the line to these problematic regions for an improved fit. **C.** Aperiodic fit (mean +/- SEM) of the power spectra in figure 3a, while the mice were in the dark with the monitor turned off (n = 44 p.g.). Dots at bottom of plots in this figure indicate points of significant difference between groups, assessed by non-parametric hierarchical bootstrap with 99% confidence. **D.** Boxplot (median, IQR, and full range) and individual data points for points of inflection (knee frequencies) from the fits in (C), WRST z-statistic = -0.672, effect size = 0.0716. **E.** Same as (D) but for the offset values (the power at 1.5 Hz) from the fits in (C), WRST z-statistic = 0.956, effect size = 0.102. **F.** Same as (D) but for the slope values of the fits in (C). WRST z-statistic = 0.705, effect size = 0.075. **G.** Bootstrapped difference (median +/- 99% CI) between the aperiodic spectra for gray screen and black screen for KO and WT mice. Red dots at the bottom indicate points of significant difference for KO, black dots for WT. **H.** Bootstrapped difference (median +/- 99% CI) between gray screen and black screen for the aperiodic offset and slope for KO and WT. **J.** Bootstrapped difference (median +/- 99% CI) between gray screen and black screen for the maximum power of Pk1a and Pk1b for KO and WT. **K.** Ratio of black screen and gray screen periodic Pk1a maximum power values for juvenile (n = 38) and adult (n = 6) KO mice, WRST z-statistic = -0.051, effect size = 0.008. **L.** Same as (K) but for periodic Pk1b maximum power values, WRST z-statistic = -2.821, effect size = 0.425. **M.** Boxplot and individual data points for the center frequency of Pk1b for juvenile (n = 67 p.g.) and adult (n = 22 p.g.) KO and WT mice viewing an iso-luminant gray screen. WRST z-statistic = 0.076, effect size = 0.007 (juvenile) and WRST z-statistic = 0.344, effect size = 0.052 (adult). **N.** Boxplot and individual data points for the center frequency of Pk1a for juvenile (n = 67) and adult (n = 22) KO mice viewing a gray screen and juvenile (n = 38) and adult (n = 6) KO mice viewing a black screen. WRST z-statistic = 4.199, effect size = 0.445 (gray screen) and WRST z-statistic = 0.706, effect size = 0.106 (black screen). **O.** Separation of juvenile KO and WT genotypes (n = 67 p.g.) by plotting Pk1b maximum power, Pk1a maximum power, and Pk1a center frequency. **P.** Separation of adult *Fmr1* KO and WT genotypes (n = 22 p.g.) by plotting Pk1a maximum power and center frequency.

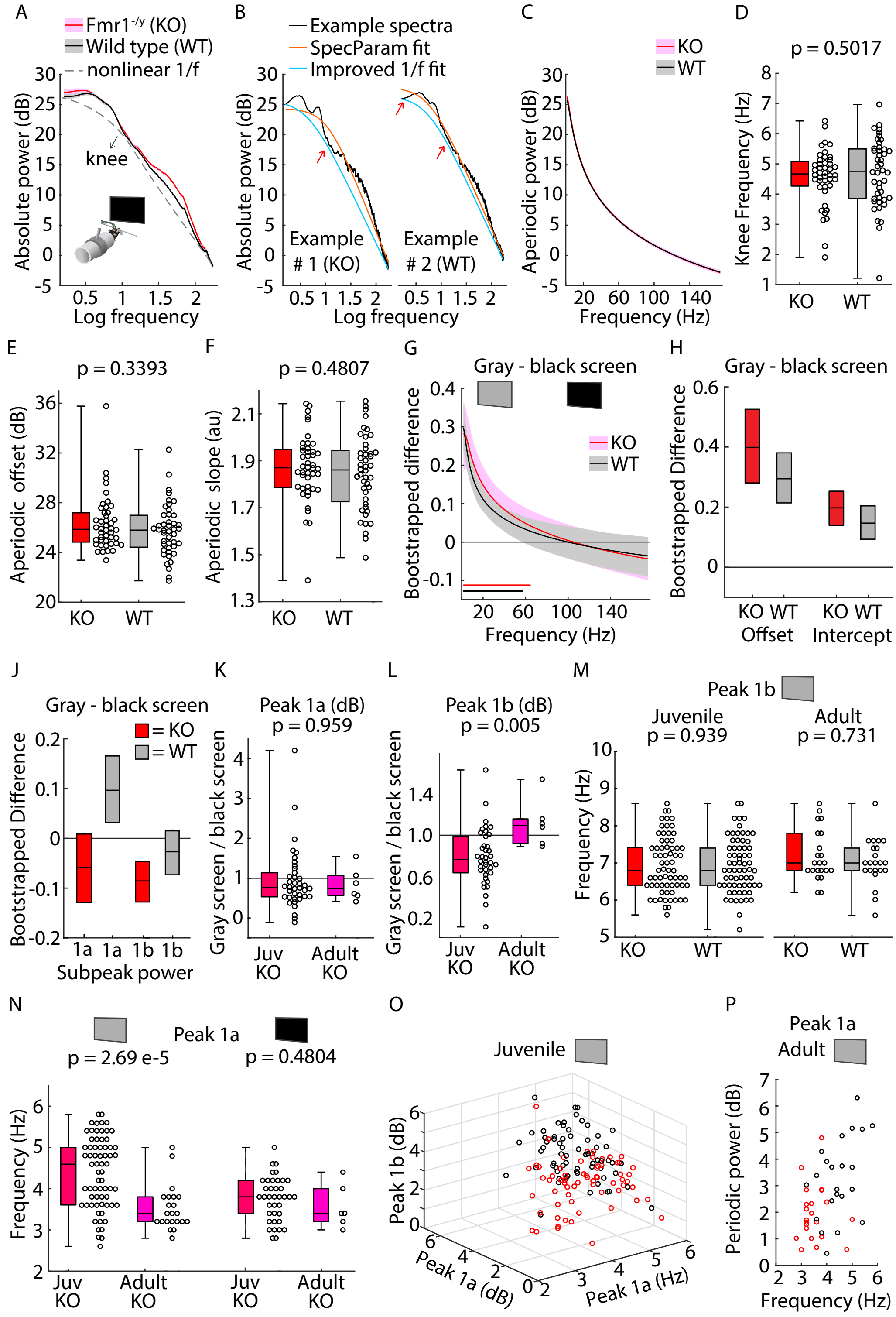

A
Fmr1-/y (KO)
Wild type (WT)
nonlinear 1/f
Absolute power (dB)
knee
Log frequency
B
Example spectra
SpecParam fit
Improved 1/f fit
Absolute power (dB)
Example # 1 (KO)
Example # 2 (WT)
Log frequency
C
KO
WT
Aperiodic power (dB)
Frequency (Hz)
D
p = 0.5017
Knee Frequency (Hz)
KO
WT
E
p = 0.3393
Aperiodic offset (dB)
KO
WT
F
p = 0.4807
Aperiodic slope (au)
KO
WT
G
Gray - black screen
Bootstrapped difference
KO
WT
Frequency (Hz)
H
Gray - black screen
Bootstrapped Difference
KO WT
Offset
KO WT
Intercept
J
Gray - black screen
= KO
= WT
Bootstrapped Difference
1a 1a 1b 1b
Subpeak power
K
Peak 1a (dB)
p = 0.959
Gray screen / black screen
Juv KO
Adult KO
L
Peak 1b (dB)
p = 0.005
Gray screen / black screen
Juv KO
Adult KO
M
Peak 1b
Juvenile
p = 0.939
Adult
p = 0.731
Frequency (Hz)
KO
WT
KO
WT
N
Peak 1a
p = 2.69 e-5
p = 0.4804
Frequency (Hz)
Juv KO
Adult KO
Juv KO
Adult KO
O
Juvenile
Peak 1b (dB)
Peak 1a (dB)
Peak 1a (Hz)
P
Peak 1a
Adult
Periodic power (dB)
Frequency (Hz)

**Extended Data 5 (corresponds to figure 3H). Cross-frequency coupling of periodic Pk1a and 14-38 Hz frequencies is impaired in *Fmr1$^{-/y}$* mice. A.** (left) Median bootstrapped noise-subtracted cross-frequency comodulogram for juvenile *Fmr1$^{-/y}$* (KO) mice (n = 67) viewing an iso-luminant gray screen showing the strength of coupling (modulation index) between the phases of periodic Pk1a oscillations (2-7 Hz) and the amplitude of higher frequency oscillations. Warmer colors indicate stronger coupling. (right) Regions of the comodulogram where the bootstrapped modulation index (MI) is significantly greater than noise (i.e., MI – noise > 0 with 99% confidence). **B.** Same as (A) but for juvenile WT mice (n = 67) viewing an iso-luminant gray screen. **C.** Probability distribution for KO mice (left) and WT mice (right) viewing a gray screen of 14-38 Hz amplitude values occurring in one of 18 bins of the periodic Pk1a (4-6 Hz) phase. A flatter distribution yields a smaller MI. **D.** Bootstrapped difference between the KO and WT mice amplitude probability distributions shown in (C). Dots at bottom of plot indicate points of significant difference between groups, assessed by non-parametric hierarchical bootstrap with 99% confidence.

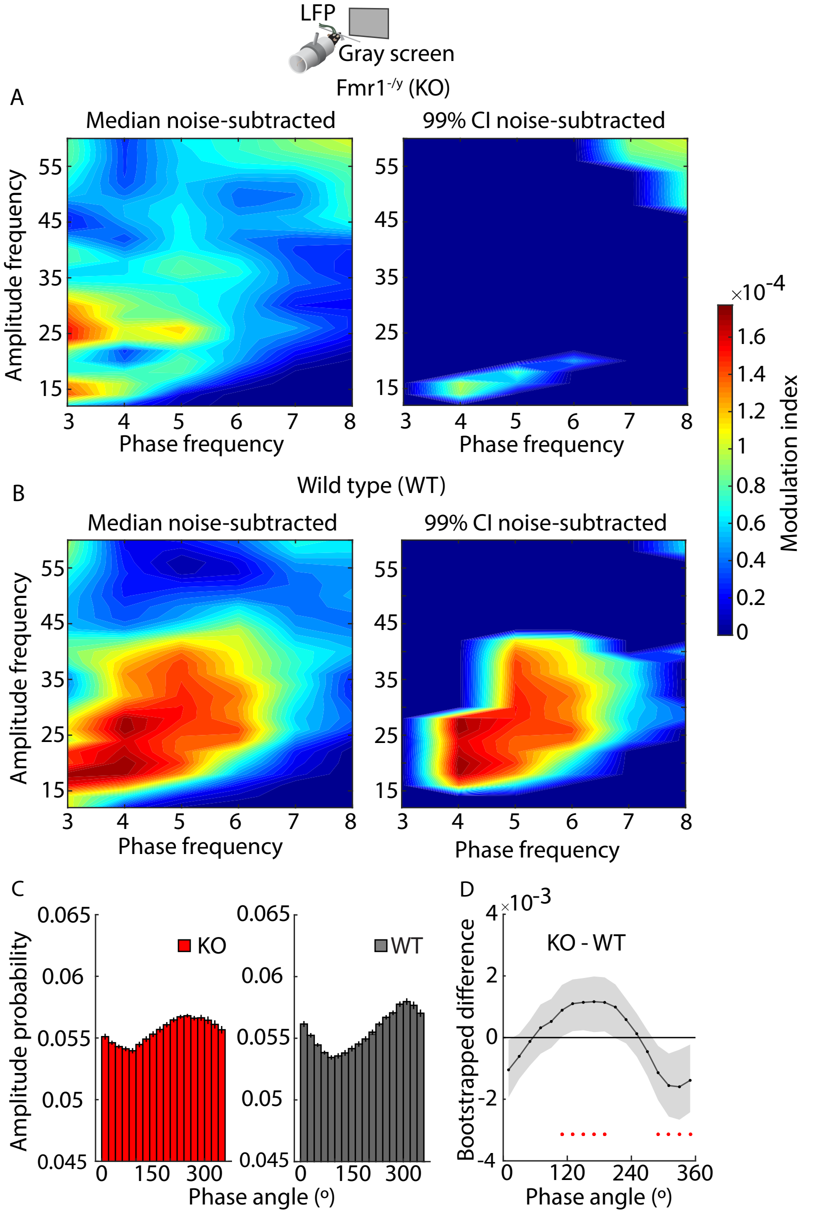

LFP
Gray screen
Fmr1-/y (KO)
A
Median noise-subtracted
99% CI noise-subtracted
Amplitude frequency
Phase frequency
×10-4
Modulation index
B
Wild type (WT)
Median noise-subtracted
99% CI noise-subtracted
Amplitude frequency
Phase frequency
C
Amplitude probability
KO
WT
Phase angle (o)
D
×10-3
KO - WT
Bootstrapped difference
Phase angle (o)

**Extended Data 6 (corresponds to figure 4). Bursting phenotypes are present in juvenile and adult *Fmr1*$^{-/y}$ mice, and altered temporal dynamics are not detectable from the surface. A.** Boxplot and individual data points for the microburst count for juvenile (left, n = 67 p.g.) and adult (right, n = 22 p.g.) KO and WT mice viewing gray screen. WRST z-statistic = 4.009, effect size = 0.386 (juvenile) and WRST z-statistic = 3.512, effect size = 0.577 (adult). **B.** Same as (A) but for microburst duration. WRST z-statistic = -4.003, effect size = 0.385 (juvenile) and WRST z-statistic = -3.511, effect size = 0.577 (adult). **C.** Distribution of average burst activity during moving and still states across all KO (left) and WT (right) animals viewing gray screen divided by juveniles (top) and adults (bottom). **D.** Percentage of time spent moving and still across all KO (left) and WT (right) animals viewing gray screen divided by juveniles (top) and adults (bottom). **E.** Comparison of burst dynamics in freely-moving KO and WT adult mice (n = 11 p.g.) with electrodes on the surface of V1 and implanted in L4 in the same hemisphere. Significant differences in microburst count (left) and duration (right) between groups were only present in L4. For L4 electrode, WRST z-statistics = -2.034 for microburst count, 2.034 for duration, effect size = 0.434 for both. For surface electrode, WRST z-statistic = +/-0.099, effect size 0.021 (for burst count and duration). **F.** Spectral dynamics across 90s of continuous data from an occipital (O2 electrode) in an example FXS (top) and TD (bottom) adult human subject. Burst activity is present in both subjects and centered around their respective periodic Pk1 maximum values (5.5-9.5 Hz for FXS, 9-12 Hz TD). **G.** Mean microburst count (left) and duration (right) across all electrodes in the occipital ROI for FXS and TD adult subject (n = 20 p.g.). WRST z-statistic = 1.041, effect size = 0.178 for counts and WRST z-statistic = -0.971, effect size = 0.1667 for durations. While group means were not significantly different, the variance of the TD distribution was significantly higher (p = 0.0018 for microburst count, p = 2.59e-5 for burst duration, Levene tests).

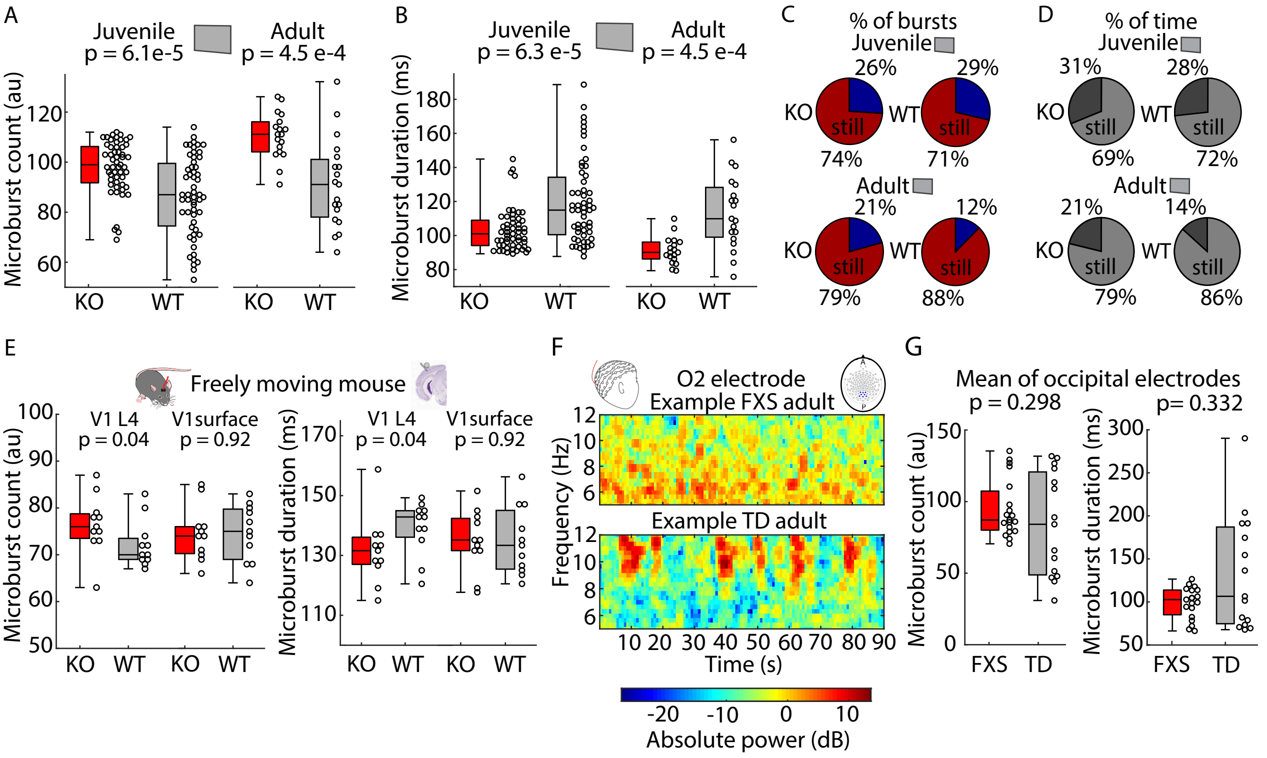
A
Juvenile
p = 6.1e-5
Adult
p = 4.5 e-4
Microburst count (au)
KO
WT
KO
WT
B
Juvenile
p = 6.3 e-5
Adult
p = 4.5 e-4
Microburst duration (ms)
KO
WT
KO
WT
C
% of bursts
Juvenile
26%
29%
KO
WT
still
74%
71%
Adult
21%
12%
79%
88%
D
% of time
Juvenile
31%
28%
69%
72%
Adult
21%
14%
79%
86%
E
Freely moving mouse
V1 L4
p = 0.04
V1surface
p = 0.92
Microburst count (au)
Microburst duration (ms)
F
O2 electrode
Example FXS adult
Example TD adult
Frequency (Hz)
Time (s)
Absolute power (dB)
G
Mean of occipital electrodes
p = 0.298
p= 0.332
Microburst count (au)
Microburst duration (ms)
FXS
TD

**Extended Data 7 (corresponds to figure 5). Detailed effects of inactivating different genetically defined classes of cortical inhibitory interneurons on the aperiodic and periodic components and temporal dynamics of the L4 LFP in V1. A.** Example fit of the aperiodic component for an example PV-Cre animal with chemogenetic actuator hM4Di expressed in V1 before (black) and after (purple) CNO systemic injection. The fit is done in log-log space and is unaffected by the notch-filtered frequencies between 45-75 Hz due to our curve-fitting algorithm. **B.** Absolute power spectrum (mean +/- SEM) from figure 5A plotted in log-log space. The mean aperiodic fit before (black) and after (purple) CNO injection in PV-Cre mice (n=16) is plotted below each corresponding absolute spectrum. **C.** Boxplot (median, IQR, and full range) and individual data points for aperiodic slope values for the fit in Fig. 5B for PV-Cre mice before and after CNO injection, WRST z-statistic = -4.805, effect size = 0.85. **D.** Same as (C) but for aperiodic knee values, WRST z-statistic = -2.695, effect size = 0.477. **E.** Boxplot and individual data points for the center frequency of periodic Pk1a and Pk1b of PV-Cre mice before and after CNO treatment. WRST z-statistic = -0.285, effect size = 0.0504 (Pk1a) and WRST z-statistic = -0.735, effect size = 0.13 (Pk1b). **F.** Aperiodic fit (mean +/- SEM) of the power spectra in figure 5K. **G** Boxplot and individual data points for aperiodic offset values (the power at 1.5 Hz) from the aperiodic fit in (E), z-statistics and effect sizes not reported for this smaller sample. **H-K.** Same as (C-E) but for SOM-Cre mice during optogenetic stimulation. **L.** Time series, 2-10 Hz band-passed signal, and piezoelectric voltage signal (as in figure 4a) but for an example PV-Cre+HM4di animal before (left) and after (right) CNO treatment. **M.** Distribution of average microburst activity during moving and still states across all PV-Cre mice before (top) and after (bottom) systemic CNO injection.

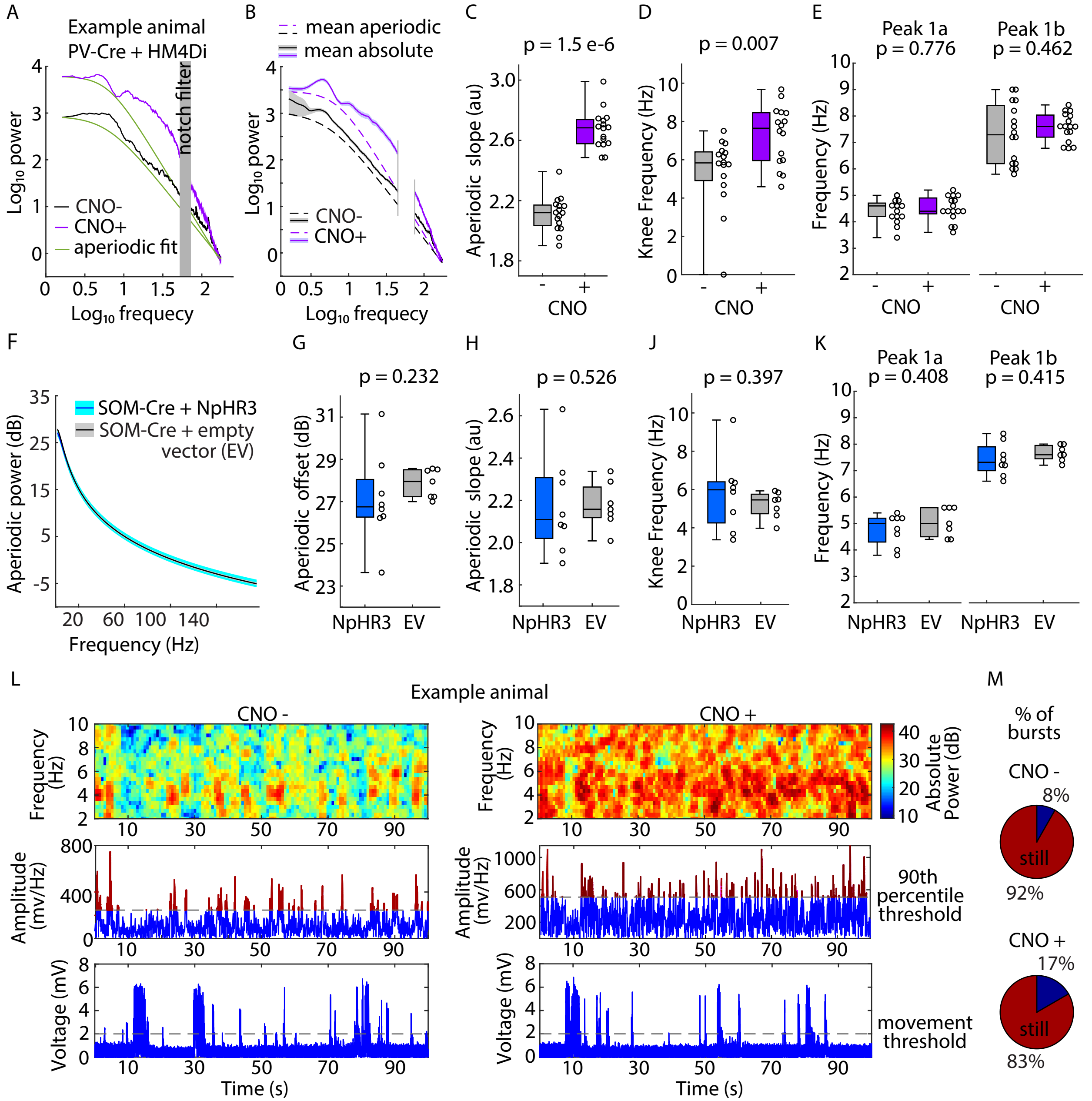

A
Example animal
PV-Cre + HM4Di
notch filter
CNO-
CNO+
aperiodic fit
Log10 power
Log10 frequecy
B
mean aperiodic
mean absolute
CNO-
CNO+
Log10 power
Log10 frequecy
C
p = 1.5 e-6
Aperiodic slope (au)
CNO
D
p = 0.007
Knee Frequency (Hz)
CNO
E
Peak 1a
p = 0.776
Peak 1b
p = 0.462
Frequency (Hz)
CNO
CNO
F
SOM-Cre + NpHR3
SOM-Cre + empty
vector (EV)
Aperiodic power (dB)
Frequency (Hz)
G
p = 0.232
Aperiodic offset (dB)
NpHR3
EV
H
p = 0.526
Aperiodic slope (au)
NpHR3
EV
J
p = 0.397
Knee Frequency (Hz)
NpHR3
EV
K
Peak 1a
p = 0.408
Peak 1b
p = 0.415
Frequency (Hz)
NpHR3
EV
NpHR3
EV
L
Example animal
CNO -
CNO +
Frequency (Hz)
Absolute Power (dB)
Amplitude (mv/Hz)
90th percentile threshold
Voltage (mV)
movement threshold
Time (s)
Time (s)
M
% of bursts
CNO -
8%
still
92%
CNO +
17%
still
83%

**Extended Data 8 (corresponds to figure 6). WT mice in the triple-transgenic experiment match expected control values, and cross-frequency coupling is impaired in EMX1-*Fmr1* KO mice. A.** Absolute power spectrum (mean +/- SEM) from the L4 LFP electrode in V1 control mice in the dark (monitor off) from the triple-transgenic EMX1-*Fmr1* KO experiment. The spectra from WT/WT (control) mice (n = 9) were compared to spectra used in figure 6A from combined Cre/WT and WT/*Fmr1* KO mice (n = 20 mice). **B.** Periodic spectra (mean +/- SEM) of the power spectra in (A). **C.** Aperiodic fits (mean +/- SEM) of the power spectra in (A). **D-E.** Microburst count (D) and duration (E) for control (WT/WT) mice vs the WT mice used in figure 6G-H. WRST z-statistic = -0.638 for count and 0.638 for duration, effect sizes = 0.125. **F.** Percentage of time spent moving across and still for 100 s across all control WT (top) and WT from figure 6J (bottom). **G.** Absolute power spectrum (mean +/- SEM) from the L4 LFP electrode in V1 of EMX1-*Fmr1* KO (Cre+/Fmr1-) and WT mice (n = 15 KO, n = 20 WT) while the mice were viewing an iso-luminant gray screen. **H.** Aperiodic fit (mean +/- SEM) of the power spectra in (G). **J.** Bootstrapped difference (median +/- 99% CI) between gray screen and black screen for the maximum power of Pk1a and Pk1b for EMX1-*Fmr1* KO and WT. **K.** Median bootstrapped noise-subtracted cross-frequency comodulogram for EMX1-*Fmr1* KO mice (left, n = 15) and WT mice (right, n = 20) viewing an iso-luminant gray screen showing the strength of coupling (modulation index) between the phases of Pk1a oscillations (2-7 Hz) and the amplitude of higher frequency oscillations. Warmer colors indicate stronger coupling. Regions of the comodulogram where the bootstrapped modulation index is significantly larger than noise with 99% confidence are outlined in black. **L.** Probability distribution for EMX1-*Fmr1* KO mice (left) and WT mice (right) viewing a gray screen of 14-38 Hz amplitude values occurring in a given phase bin, where the phase of periodic Pk1a oscillations (4-6 Hz) was divided into 18 bins. A flatter distribution yields a smaller MI. **M.** Bootstrapped difference between the EMX1-*Fmr1* KO and WT mice amplitude probability distributions shown in (C).

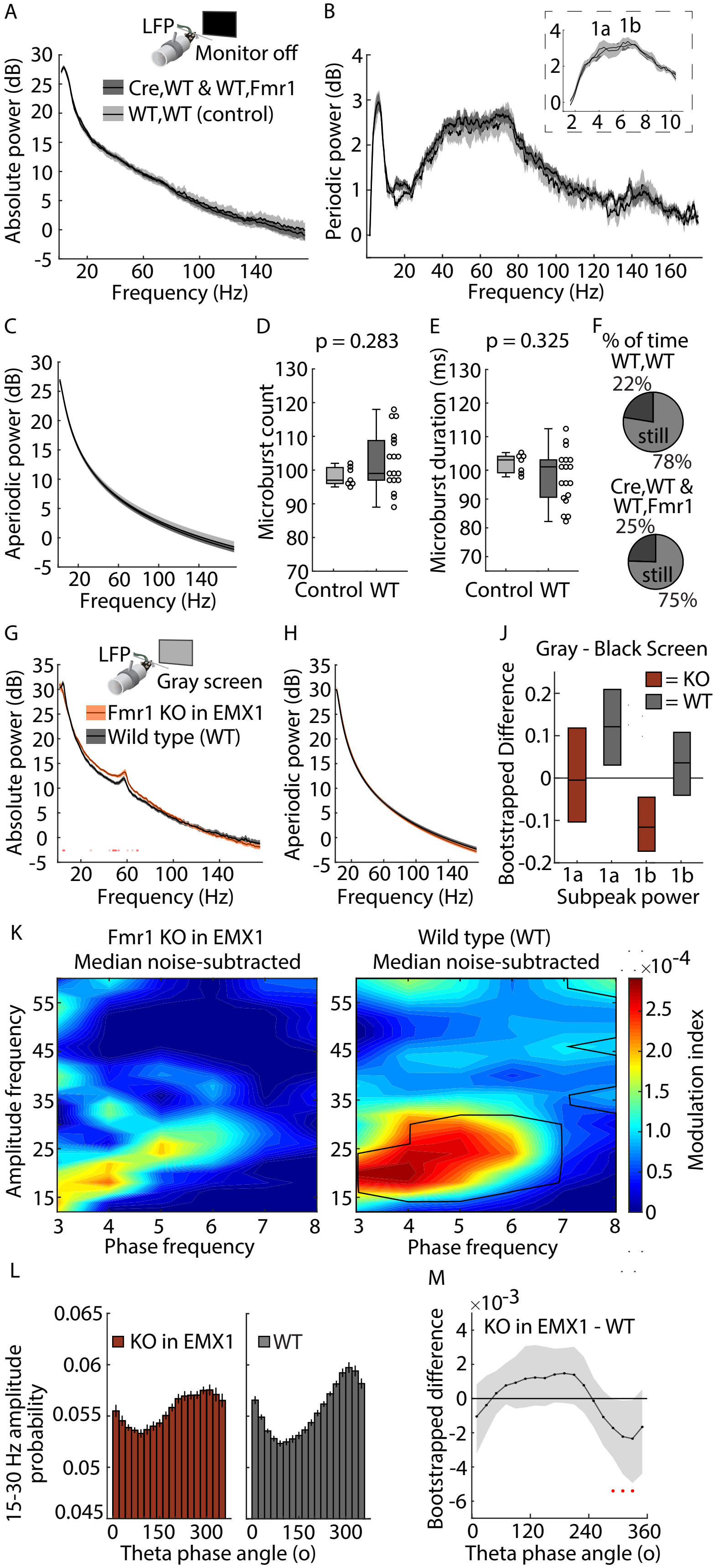

A
LFP
Monitor off
Cre,WT & WT,Fmr1
WT,WT (control)
Absolute power (dB)
Frequency (Hz)
B
1a 1b
Periodic power (dB)
Frequency (Hz)
C
Aperiodic power (dB)
Frequency (Hz)
D
p = 0.283
Microburst count
Control WT
E
p = 0.325
Microburst duration (ms)
Control WT
F
% of time
WT,WT
22%
still
78%
Cre,WT &
WT,Fmr1
25%
still
75%
G
LFP
Gray screen
Fmr1 KO in EMX1
Wild type (WT)
Absolute power (dB)
Frequency (Hz)
H
Aperiodic power (dB)
Frequency (Hz)
J
Gray - Black Screen
= KO
= WT
Bootstrapped Difference
1a 1a 1b 1b
Subpeak power
K
Fmr1 KO in EMX1
Median noise-subtracted
Wild type (WT)
Median noise-subtracted
Amplitude frequency
Phase frequency
×10^-4
Modulation index
L
KO in EMX1
WT
15-30 Hz amplitude probability
Theta phase angle (o)
M
×10^-3
KO in EMX1 - WT
Bootstrapped difference
Theta phase angle (o)

**Extended Data 9 (corresponds to figure 7C-E). Detailed genotype differences in VEPs for stimulus onset and offset. A.** Probability distribution for WT mice (left, n = 31) and *Fmr1*$^{-/y}$ (KO) mice (right, n = 31) of the phase of the alpha-like oscillations at the time of stimulus onset occurring in one of 18 phase bins. **B.** Bootstrapped difference between the KO and WT mice phase probability distributions shown in (A). Dots at bottom of plots in this figure indicate points of significant difference between groups, assessed by non-parametric hierarchical bootstrap with 99% confidence. **C.** In both EMX1-*Fmr1* KO (n = 15) and littermate WT mice (n = 29), N1 magnitude was correlated with the phase of the band-passed 2-10 Hz signal at the time of stimulus onset. **D.** Onst response broken down by temporal frequency of the stimulus' phase reversals for KO and WT mice. While the positivity around 125 msec varied across stimuli, the initial negativity (~75 msec) was stereotyped across all frequencies. **E.** Onset response for juvenile (left, p30-40) and adult (right, p70-100) KO and WT mice. Genotype differences persisted across development. **F.** Onset response across all KO and WT mice separated by the movement state of the animal during the 250 msec period. Genotype differences persisted during movement. **G.** Matching distribution of onset responses across all KO and WT mice that occurred during movement bouts (left) and still bouts (right). The majority of onset responses occurred while the mice were not moving. **H.** There was also a stereotyped response when the grating stimuli ended and gray screen resumed (offset response). As previously reported, this response had about 3 oscillations[73]. Here we show it was reduced in magnitude in KO mice relative to WT, even after unfamiliar stimuli. **H.** There was no difference between EXM1-*Fmr1* KO and littermate WT mice in the amount of forepaw movement in the three seconds after stimulus onset.

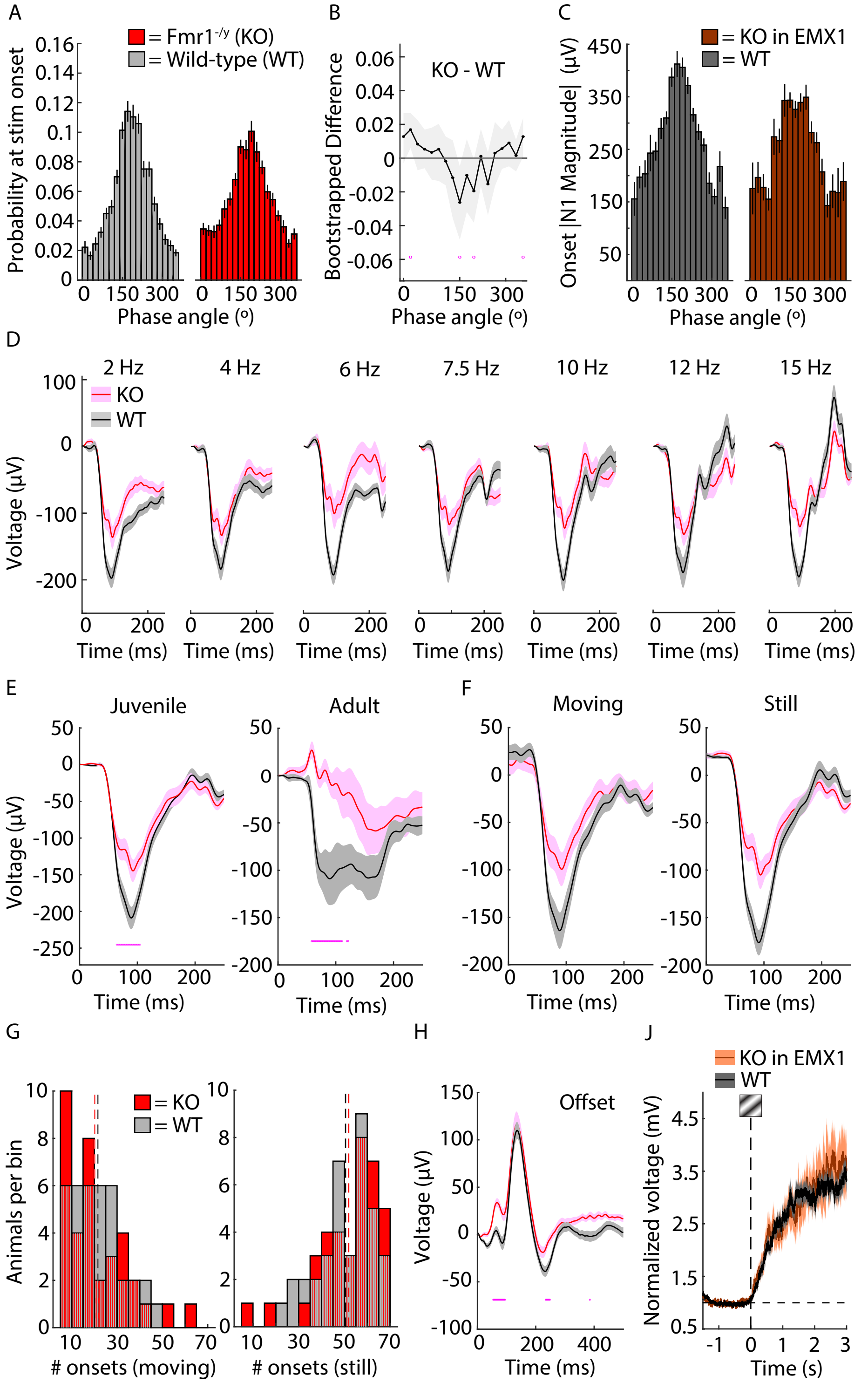
A
= Fmr1-/y (KO)
= Wild-type (WT)
Probability at stim onset
Phase angle (o)
B
KO - WT
Bootstrapped Difference
Phase angle (o)
C
= KO in EMX1
= WT
Onset |N1 Magnitude| (µV)
Phase angle (o)
D
2 Hz
4 Hz
6 Hz
7.5 Hz
10 Hz
12 Hz
15 Hz
KO
WT
Voltage (µV)
Time (ms)
E
Juvenile
Adult
Voltage (µV)
Time (ms)
F
Moving
Still
Time (ms)
G
= KO
= WT
Animals per bin
### onsets (moving)
### onsets (still)
H
Offset
Voltage (µV)
Time (ms)
J
KO in EMX1
WT
Normalized voltage (mV)
Time (s)

**Extended Data 10 (corresponds to figure 7f-g). ssVEP results are consistent with analysis in the time domain. A.** 1- or 2-sec segments of LFP data during “entrainment” periods represented in the frequency domain (mean +/- SEM) for each stimulation frequency. ssVEP amplitudes were the value at the stimulation frequencies (marked with blue arrows). Dots at bottom of plots in this figure indicate points of significant difference between groups, assessed by non-parametric hierarchical bootstrap, and blue dots indicate a significant difference at the stimulation frequency with 99% confidence. **B.** ssVEP amplitudes normalized to the ssVEP magnitude during equivalent durations of static gray screen for each stimulation frequency to remove the contribution of resting-state Pk1 periodic power. * indicate temporal frequencies for which there is a significant difference between genotypes assessed through nonparametric hierarchical bootstrapping with 99% confidence. **C.** Average LFP data aligned to each phase reversal during “entrainment” periods represented in the time domain (mean +/- SEM) for the corresponding duration of each stimulation frequency. A complete (two-peak) VEP waveform cannot be ascertained above 10 Hz. **D.** Bootstrapped difference (median +/- 99% CI) between genotypes the peak-to-peak amplitude of the VEP for each temporal frequency and broken down by age (n = 31 juvenile, n = 6 adult). Significant differences are found where the confidence interval does not overlap with 0. **E.** Percentage of time spent moving and still across all *Fmr1* KO (top) and WT (bottom) animals during all periods of grating presentations. **F.** Same as (B) but for EMX1-*Fmr1* KO (n = 15) and littermate WT (n = 29) mice. **G.** Same as (E) but for EMX1-*Fmr1* KO and WT mice. **H-J.** Same as (C-D) but for EMX1-*Fmr1* KO and WT mice.

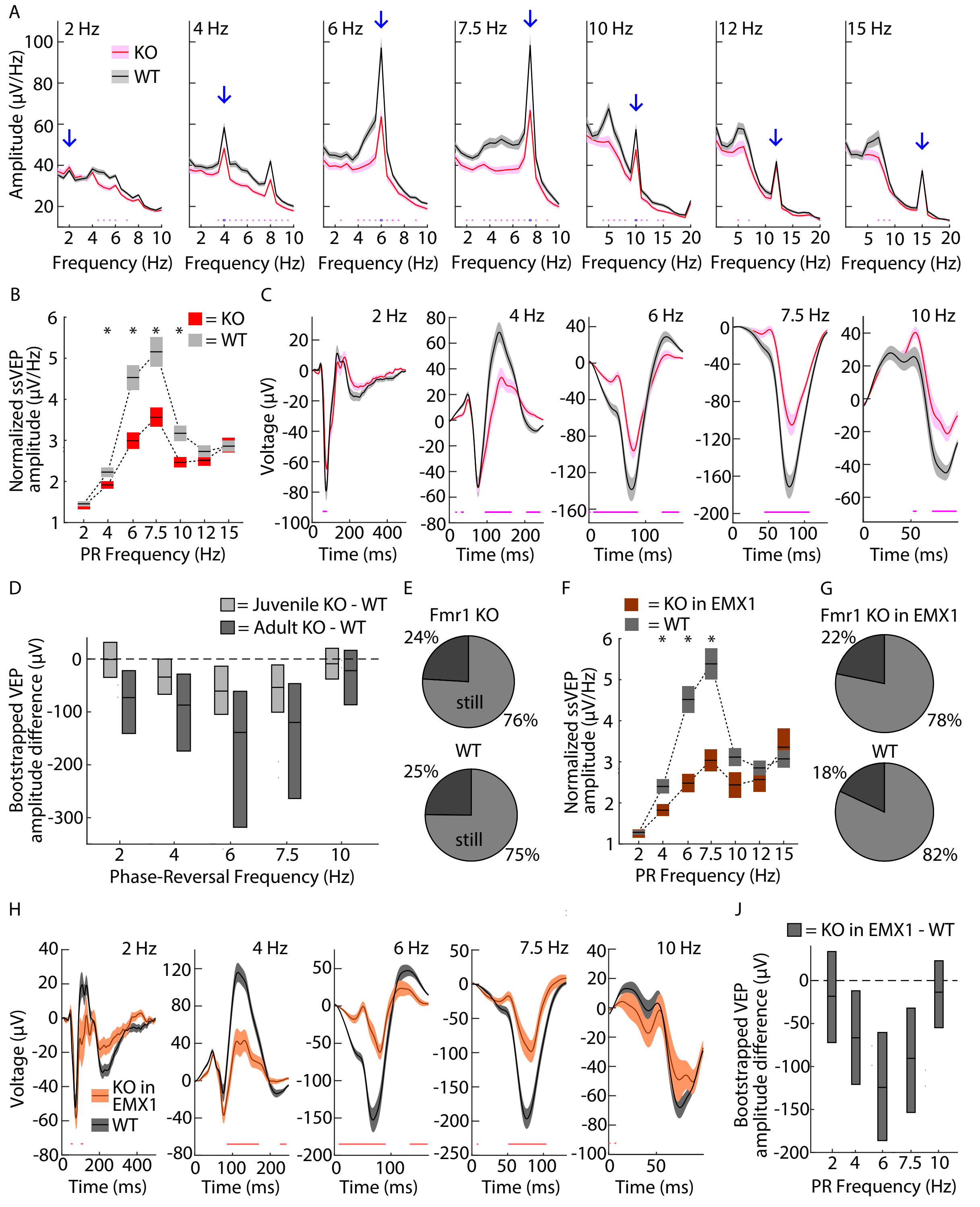

A
2 Hz
4 Hz
6 Hz
7.5 Hz
10 Hz
12 Hz
15 Hz
KO
WT
Amplitude (µV/Hz)
Frequency (Hz)
B
= KO
= WT
Normalized ssVEP amplitude (µV/Hz)
PR Frequency (Hz)
C
Voltage (µV)
Time (ms)
D
= Juvenile KO - WT
= Adult KO - WT
Bootstrapped VEP amplitude difference (µV)
Phase-Reversal Frequency (Hz)
E
Fmr1 KO
24%
still
76%
WT
25%
still
75%
F
= KO in EMX1
= WT
Normalized ssVEP amplitude (µV/Hz)
PR Frequency (Hz)
G
Fmr1 KO in EMX1
22%
78%
WT
18%
82%
H
KO in EMX1
WT
Voltage (µV)
Time (ms)
J
= KO in EMX1 - WT
Bootstrapped VEP amplitude difference (µV)
PR Frequency (Hz)
